# Layer-specific cortical processing dissociates sensory and cognitive influences on pain

**DOI:** 10.64898/2026.05.12.724526

**Authors:** Balint Kincses, Viktor Pfaffenrot, Katharina Püchner, Tamas Spisak, Katja Wiech, Peter Koopmans, Ulrike Bingel

## Abstract

Pain arises from the integration of nociceptive input with cognitive and affective processes, yet how these signals are organized within cortical circuits remains unclear. Here, using submillimeter-resolution 7T fMRI, we tested whether bottom-up (BU) and top-down (TD) influences on pain are segregated across cortical layers in human sensory-discriminative regions. Participants underwent a factorial manipulation of nociceptive input and cognitive modulation, enabling dissociation of BU and TD processes. BU processing was strongest in middle cortical layers. In contrast, TD modulation preferentially engaged superficial layers, consistent with cortico-cortical feedback mechanisms. Critically, individual differences in this laminar segregation predicted the magnitude of distraction-induced analgesia, linking circuit-level organization to behavior.

These findings provide evidence that cognitive modulation of pain is implemented through layer-specific cortical computations and extend canonical microcircuit models to human pain processing. More generally, they establish laminar fMRI as a powerful approach for linking cortical circuit architecture to subjective experience.

## Introduction

Pain is a complex experience that extends beyond the mere feedforward transmission of nociceptive input. In addition to bottom-up (BU) sensory signals, pain perception is profoundly shaped by cognitive and affective processes such as expectation and attention^1,2^. Identifying the neurobiological underpinnings of these processes, particularly at the level of the neocortex, would fundamentally advance our understanding of pain mechanisms.

Traditional models of pain processing have mapped nociceptive input onto early cortical pain regions, most prominently the primary and secondary somatosensory cortices (S1 and S2) and the posterior insula (pIns)^3–5^. These regions have therefore often been conceptualized as primary sensory or nociceptive areas. However, accumulating evidence shows that activity within these areas is also modulated by cognitive factors^6–8^, suggesting that they enable the parallel processing of BU and top-down (TD) signals, rather than acting as passive recipients of nociceptive input.

The canonical cortical microcircuit model^9^, a hierarchical model of cortical processing, provides a mechanistic framework to probe the neural basis of the overlapping representation of these processes. This model assigns different functions to specific cortical layers^10,11^, whereby feedforward signals (BU) predominantly target the middle (granular) layers, whereas feedback signals (TD) target superficial and deep (agranular) layers^12^. Applied to pain, this framework suggests that nociceptive and cognitive signals are segregated across cortical layers. This hypothesis has long remained difficult to test directly, as conventional neuroimaging methods lacked the spatial resolution to resolve laminar-specific activity. Advances in submillimeter-resolution fMRI now make it possible to investigate these fine-grained cortical processes in humans^13–15^. Using these methods, studies in the visual^10,13^, auditory^16^, and somatosensory^17^ systems have already demonstrated laminar dissociations between BU and TD processing. Despite these advances, progress in understanding the laminar level representation of pain-related processes remains limited.

Here we investigated whether the cortical representation of sensory and cognitive modulation of pain is implemented through laminar-level processing. In this pre-registered study, we hypothesized that BU nociceptive input preferentially engages the middle cortical layers, whereas TD cognitive modulation is predominantly expressed in agranular (superficial and deep layers (https://osf.io/68je5). To test this hypothesis, we independently manipulated nociceptive input and cognitive load within a well-established experimental paradigm while acquiring submillimeter-resolution fMRI data at ultra-high field. By resolving pain-related activity across cortical depths, this study directly tests a key prediction of hierarchical models of cortical pain processing, providing mechanistic insight into how sensory and cognitive processes are represented in the neocortex.

## Results

To investigate the sensory and cognitive modulation of pain perception, we applied a well-established within-subject 2 by 2 factorial design^18^ to modulate BU and TD effects of pain. The cognitive modulation consisted of a modified version of an n-back task with two different levels of cognitive demand (*low demand* condition (0-back) and *high demand* condition (2-back), *Supplementary material - Task and stimuli*) which were combined with concomitant low or high intensity thermal stimulation (factor temperature; levels: high and low; see Figure 1 and Supplementary Figure 1). Performance of an n-back task during noxious stimulation has been shown to reduce pain intensity ratings with a stronger *distraction* effect the more difficult the cognitive task is^19,20^. Because our study focuses on the modulatory effect of cognitive demand, we will refer to the first factor as distraction (weak distraction in the 0-back condition vs. strong distraction in the 2-back condition). The main effect of distraction reflecting TD is therefore defined as ‘weak minus strong distraction’ and the main effect of temperature reflecting BU is defined as ‘high minus low temperature’. Because our study focuses on the investigation of TD modulatory processes induced by a cognitive task (as opposed to the question of whether the cognitive task modulates pain intensity ratings at all), we investigated a sample of individuals (N = 30) who passed a preregistered behavioral threshold, thereby ensuring that only participants who showed significantly reduced pain ratings during the performance of the strong distraction task were included (relative to the weak distraction task) (see *Methods* and Supplementary Figure 1 for more detail; https://osf.io/68je5). As expected, the behavioral results replicated each modulatory effects in-scanner, showing a main effect of temperature with higher pain intensity ratings at higher stimulus temperature (ES: 33.23 VAS100, CI [30.98, 35.49], p<0.001, Figure 1B) as well as successful cognitive modulation, with higher pain ratings in the weak distraction task as compared to the strong distraction task (main effect of distraction) (effect size (ES): 6.6 VAS100, confidence interval (CI): [4.34, 8.86], p<0.001, Figure 1B). Additionally, we found a significant interaction effect (ES: 3.52, CI [0.32, 6.71], p=0.031), with a stronger distraction effect in the high temperature condition (Figure 1B, Supplementary Figure 2).

**Figure 1.**
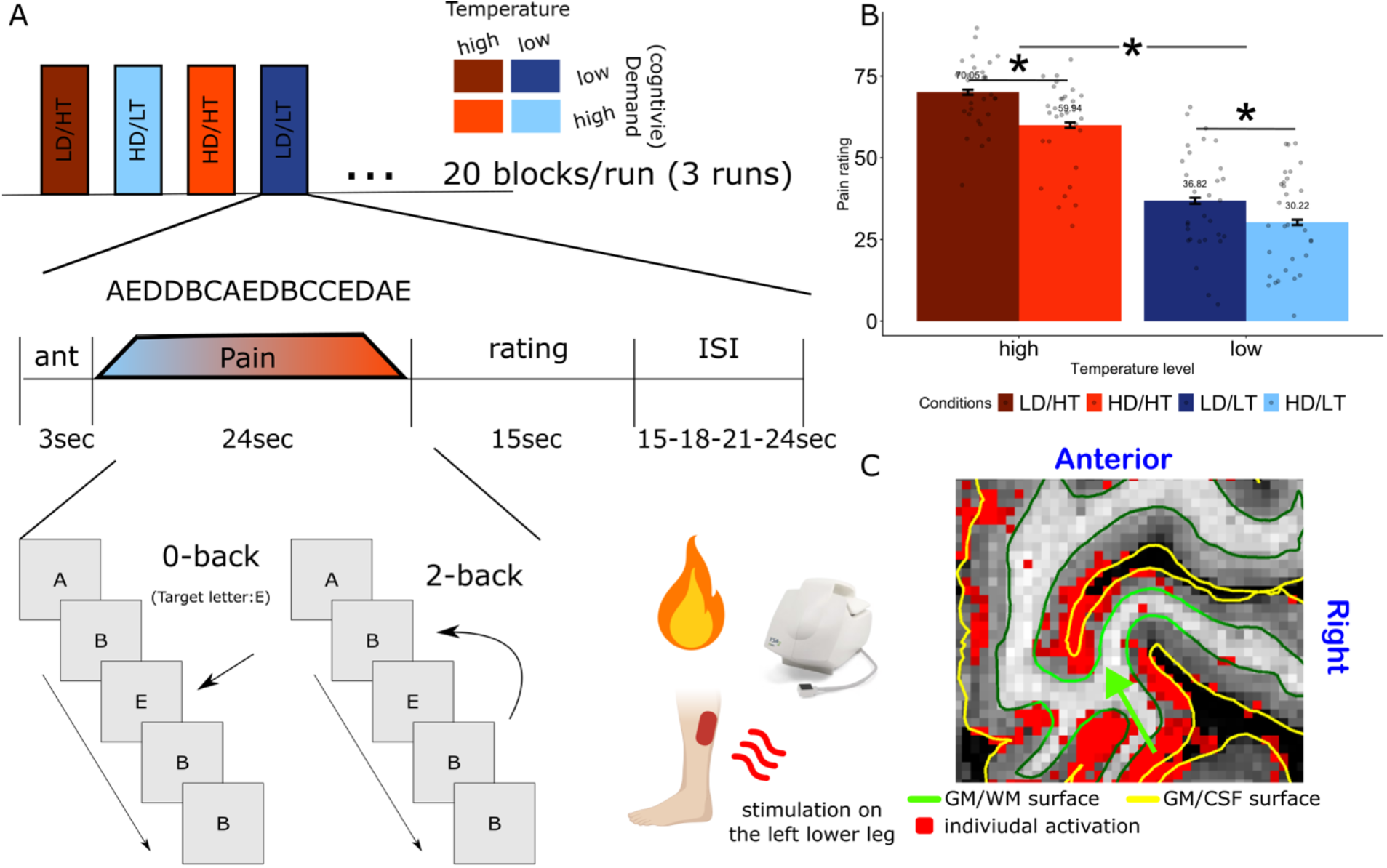
Experimental paradigm, behavioral effects, and example laminar region of interest. A. Experimental paradigm and task. The experimental design and its depiction were adopted from Sprenger et al.^18^. Participants performed an n-back task with two different levels of cognitive demand (low demand: 0-back; high demand: 2-back condition). These conditions reflect the weak and strong distraction levels, respectively. A concurrent painful heat stimulation was delivered on the participant’s left leg. Two individually calibrated temperature levels were applied (low and high nociceptive input, corresponding to low and high perceived pain). Each trial began with a 3-s anticipation period presenting the upcoming task instruction, followed by a 24-s task period and a rating phase. B. Behavioral results demonstrate robust top-down modulation by cognitive demand (main effect of distraction; ES = 6.6, CI = [4.34, 8.86], p < 0.001) and a strong bottom-up effect of nociceptive input (main effect of temperature; ES = 33.23, CI = [30.98, 35.49], p < 0.001; n = 30; see Supplementary Fig. 2 for the full sample). The significant interaction (ES = 3.52, CI = [0.32, 6.71], p = 0.031) indicates that cognitive modulation of pain is more pronounced at higher levels of nociceptive input. Asterisks denote p < 0.05; error bars indicate within-subject standard error. LD = low demand; HD = high demand; LT = low temperature; HT = high temperature. C. Example single-participant functional region of interest in primary somatosensory cortex (S1), showing activation associated with increased nociceptive input overlaid on a high-resolution anatomical image. Activation maps were anatomically smoothed (Laynii: LN_GRADSMOOTH^21^). High resolution anatomical image is in the background. Green indicates the white-matter/gray-matter boundary; yellow indicates the pial surface. The lime arrow and surface indicate the individually defined S1 region of interest projected onto the white-matter/gray-matter boundary.

High resolution functional magnetic resonance images were acquired on a 7T scanner (MAGNETOM Terra, Siemens Healthcare, Erlangen, Germany). We focused our analysis on the investigation of the laminar profiles of the primary (S1) and secondary somatosensory (S2) cortices and the posterior insula (pIns) (Supplementary Figure 3). These regions are core components of the nociceptive system and exhibit a well-defined granular laminar organization, providing an anatomically grounded framework for examining laminar-specific activity patterns. Individually defined functional ROIs were created by intersecting group-level functional activity maps with preregistered anatomical regions and restricting them to individually responsive areas showing both effects (Figure 1C and see *Methods* – *Functional ROI definition* and *Sensitivity analysis* for the robustness of the selection procedure).

Differences in laminar patterns between the two processes (main effects of temperature, BU, and main effect of distraction, TD) were compared following the analysis approach used by Lawrence et al^15^.

The main effect of temperature, reflecting BU processing, showed a clear inverted U-shape laminar profile, with strongest activation in the middle cortical layers in all regions-of-interest (Figure 2A, Table 1, https://github.com/kincsesbalint/LaminarfMRI_inPainPerception). Modeling this laminar profile with a second-order polynomial model revealed a significant negative quadratic term across all investigated regions (for left S2 data see Table 1; quadratic coefficient in BU process: b = -6.24, SE = 0.29, t(1192) = -21.63, p < .001, 95% CI [-6.80, - 5.67]; for all other regions see Supplementary Table 2-7, Supplementary results – Sensitivity analysis).

**Figure 2.**
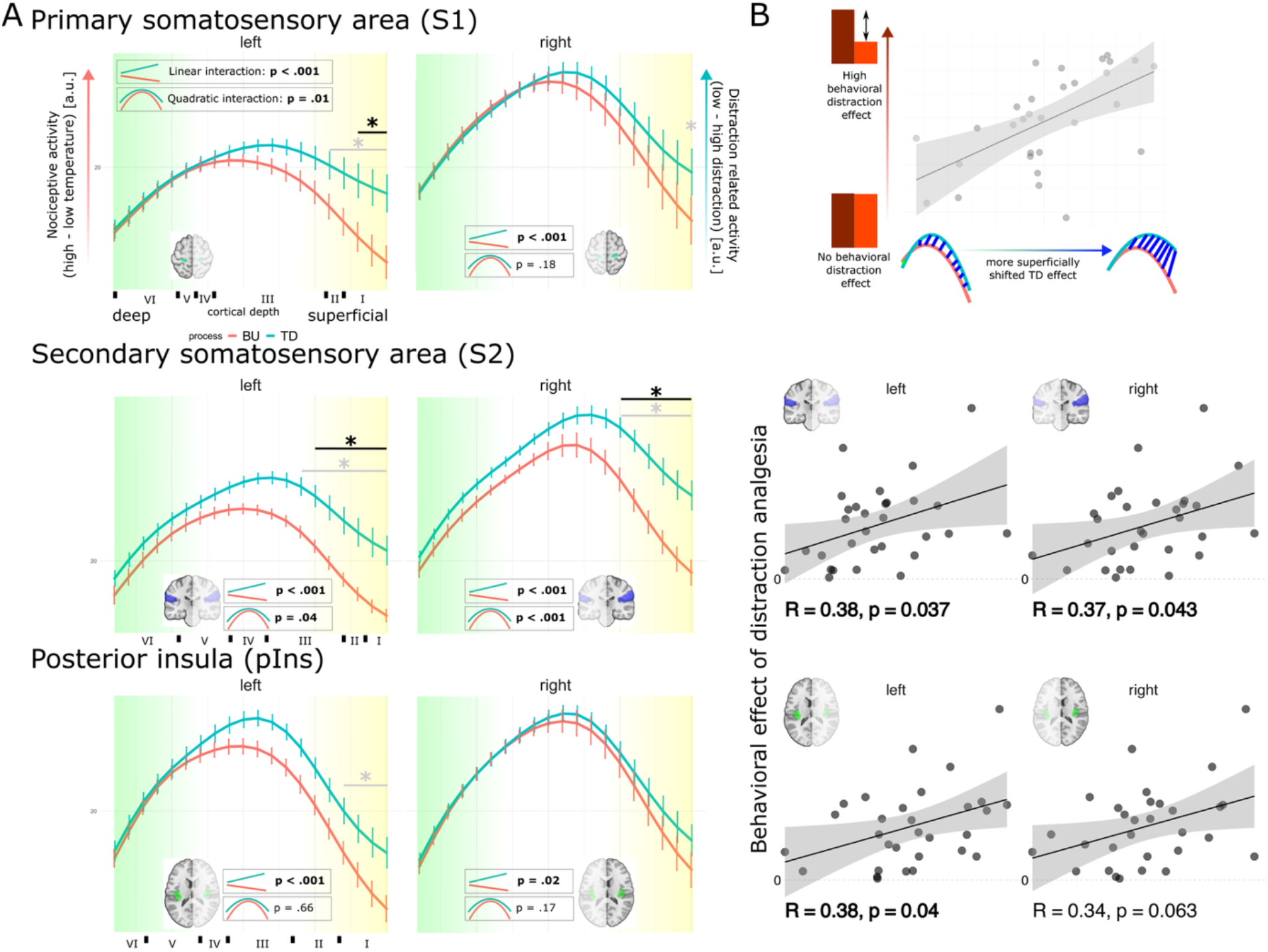
Laminar dissociation of bottom-up and top-down pain processing and its behavioral relevance. A. Laminar profiles of bottom-up (BU) and top-down (TD) effects are shown for left and right hemispheres across sensory-discriminative regions of interest (S1, S2, and posterior insula [pIns]). BU effects were defined as the main effect of nociceptive input (high–low temperature), and TD effects as the main effect of distraction (weak–strong distraction). Linear mixed-effects models were fitted separately for each hemisphere, with process (BU vs TD) and cortical layer as fixed effects. Given the inverted U-shaped laminar profiles, a second-degree polynomial model with linear and quadratic terms was fitted, and the terms’ interactions with process were tested (see Supplementary Results—Model fitting). A significant process × layer interaction indicates distinct laminar dependencies for BU and TD effects. Statistics for linear and quadratic interaction terms are shown in black boxes; significant effects are highlighted in bold. To localize the layers driving these interactions, follow-up models were performed. Black asterisks indicate FDR-corrected significant effects at the corresponding layer (p < 0.05); gray asterisks indicate significant uncorrected effects. Color coding follows Fig. 1, with green denoting deeper layers (near the GM/WM boundary) and yellow denoting superficial layers (near the GM/CSF boundary); n = 30. Laminar depth distributions (x-axis) are based on histological data from Palomero-Gallagher & Zilles^22^ for S1, Eickhoff et al.^23^ for S2, and Kurth et al.^24^ for posterior insula (Ig1 subregion). Points indicate the mean, and error bars represent the standard error of the mean. B. Relationship between laminar patterns and behavioral analgesia. Behavioral analgesia (y-axis) is plotted against individual differences in relative agranular involvement (x-axis) (see Statistical analysis). Higher values indicate greater agranular involvement of TD relative to BU process. The upper panel shows synthetic data for illustration. Significant associations were observed in S2 and pIns, indicating that individuals with a stronger agranular involvement of TD effects is associated with greater behavioral analgesia. No significant association was observed in S1 (see Supplementary Fig. 4). Correlations were assessed using two-sided Spearman tests.

**Table 1.**
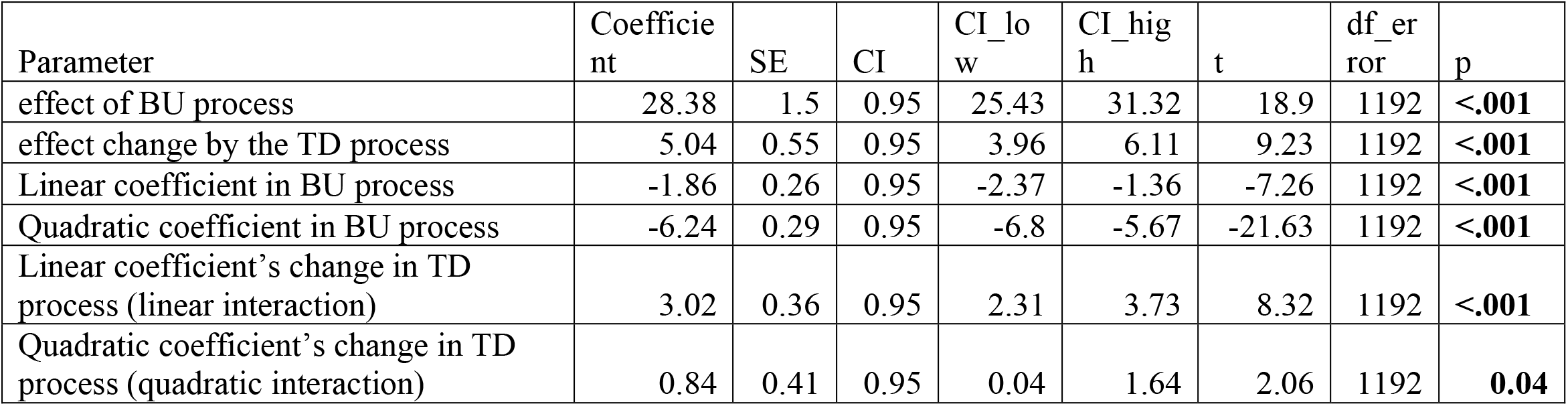
Parameter estimates from the mixed-effects model fitted to left S2. The model reveals a significant main effect of the top-down (TD) process, indicating greater activation during cognitive modulation compared with bottom-up (BU) processing. Linear and quadratic terms jointly describe the laminar profile shape (see Supplementary Methods): a negative quadratic coefficient indicates an inverted U-shaped profile peaking in middle layers, while the linear coefficient determines whether this peak is shifted toward deeper layers (negative values) or superficial layers (positive values). Importantly, the interaction with process indicates a change in laminar signaling for TD relative to BU process, such that the linear coefficient reverses sign (−1.86 + 3.02 = +1.16) and the quadratic term becomes less negative (−6.24 + 0.84 = −5.40), consistent with a relative shift toward superficial layers during TD modulation. Significant interaction terms (p < 0.05) indicate that the laminar profile of the TD process differs reliably from that of the BU process. Follow-up modeling (see Methods—Statistical analysis) directly compared these effects at each layer and identified superficial layers as primary contributors to this difference (Fig. 2A; Supplementary Tables 8–13).

In contrast to the laminar profile of BU process, TD processing showed a preferential engagement of superficial layers across S1, S2, and posterior insula, as reflected in a significant statistical interaction indicating a shift in activation along the cortical depth axis (from deep to superficial layers; captured by the linear term of our second-order polynomial regression approach (see boxes in Figure 2A) ) and in the relative degree of middle-layer-centered activation (captured by the quadratic term (Figure 2A, Table 1, Supplementary Table 2-7). For instance, in left S2, model coefficients indicated a superficial shift in the TD effect. The linear term’s coefficient estimate reversed sign for TD processing (b_linear_: +1.14 (TD) vs -1.86 (BU); b_Δlinear in TD_ = +3.02; SE = 0.36, t(1192) = 8.32, p < .001, 95% CI [2.31, 3.73]), indicating increased relative activation in superficial layers for TD relative to BU processing. Additionally, the quadratic term’s coefficient estimate is less negative (b_quadratic_: -5.4 (TD) vs - 6.24 (BU); b_Δquadratic in TD_ = + 0.84, SE = 0.41, t(1192) = 2.06, p = .040, 95% CI [0.04, 1.64]) in the TD process, indicating a flatter inverted U-shape profile and higher relative contribution of superficial layers. Together, these terms indicate a flatter inverted U-shape profile with a peak shifted toward superficial layers in the TD process. This pattern was robust across analytical choices, including region-of-interest definition (*Methods – Sensitivity analysis*, Supplementary code, Supplementary Figure 8-9).

To identify which layers contributed most strongly to the observed differences in laminar profiles between BU and TD processing, we conducted a more flexible (but less well powered) modeling approach in which layer was treated as a discrete categorical variable (see *Methods - Statistical analysis* for more detail). In left-sided cortical regions (ipsilateral to the stimulation site), the significant interaction effect between ‘layer’ (from deep to superficial) and ‘process’ (TD vs. BU) was observed in the superficial layers (Figure 2A, FDR corrected p<0.05, Supplementary Table 8-13, https://github.com/kincsesbalint/LaminarfMRI_inPainPerception). On the right side (contralateral to the stimulation), this superficial layer contribution was observed only in S2 (FDR corrected p<0.05). In summary, laminar activation profiles differed between TD and BU processing across all investigated cortical regions, with TD effects showing a more superficial activation profile than BU effects.

To investigate the functional relevance of the laminar differences between TD and BU processing, we examined their relationship with behavioral analgesia (Figure 2B). We quantified the difference between the two laminar activation profiles as an individual relative agranular involvement (see *Methods – Statistical analysis* for more detail), that captures the extent to which relative activation was shifted toward agranular layers during TD relative to BU processing (note that “shift” refers to a relative difference in laminar weighting between TD and BU processing and does not imply a temporal change in activation across cortical depth). This measure was derived from subject-specific estimates obtained from the mixed-effects model (see *Methods – Statistical analysis* for more detail). This analysis revealed that a greater agranular involvement of the TD effect was associated with stronger behavioral analgesia in S2 (left: r=0.38, p=0.037; right: r=0.37, p=0.043, Figure 2B) and posterior insula (left: r=0.38, p=0.04; right: r=0.34, p=0.063, Figure 2B), but not in S1 (left: r=0.18, p=0.33; right: r=0.23, p=0.21, Supplementary Figure 4). These findings support the behavioral relevance of the observed laminar difference.

## Discussion

In this high-resolution 7T fMRI study, we observed distinct laminar profiles of bottom-up and top-down processes of pain-related brain activity in cortical pain regions, including the primary and secondary somatosensory cortices and the posterior insula. Specifically, these laminar profiles suggest that superficial layers are differentially engaged by TD and BU processes. Notably, we found that individual differences in laminar organization predicted the magnitude of distraction-induced analgesia. Greater relative agranular engagement during TD processes was associated with stronger distraction-induced analgesia. These findings demonstrate the feasibility of applying high-resolution fMRI to pain research.

Cortical mechanisms have long been a central focus in pain neuroscience, yet empirical evidence from human studies remains sparse. Although theoretical^25^ and computational models^26^ have proposed that cognitive modulation operates via feedback signals targeting agranular cortical layers, this prediction has not been directly tested in humans. By resolving cortical activity across layers in sensory-discriminative pain regions with ultra-high field fMRI, our study provides evidence that cognitive modulation of pain is implemented through a differential weighting of superficial layer activity, rather than uniform attenuation of nociceptive signals. These results suggest that cognitive modulation of pain preferentially engages cortico-cortical feedback mechanisms, while remaining compatible with parallel modulation of bottom-up input. This extends the canonical microcircuit framework to pain and demonstrates that principles previously established in other sensory domains^10,13,16,17,27^ generalize to pain perception. Together, these results establish a circuit-level account of cognitive pain modulation in humans and identify depth-specific processing as a key mechanism linking cognition to subjective pain experience.

The functional relevance of laminar segregation is underscored by its direct relationship to behavior, demonstrating that the observed laminar pattern is not merely a structural feature but mechanistically linked to perception. Our results move beyond regional descriptions of pain intensity coding^28–31^ by uncovering a layer-specific circuit mechanism in which cortical contributions depend on the differentiation of BU and TD signals across cortical layers. This association suggests that effective pain modulation relies on increased segregation of sensory and cognitive inputs within the local cortical circuitry of these regions. Together, these results position sensory-discriminative cortices as active computational sites where laminar-specific processing shapes the influence of cognitive context on perceived pain.

The observed segregation of BU and TD signals does not imply independent processing streams, but rather support their recurrent integration within local cortical microcircuits. According to the canonical microcircuit model, signals are continuously exchanged across layers via vertical connections, allowing sensory and contextual information to be integrated throughout the cortical column^25,32^. Within this framework, cognitive modulation is expected to be most prominent in agranular layers, which are the principal targets of cortico-cortical feedback and the main source of forward projections^12,25,32^. Our findings are consistent with this architecture: although nociceptive signals remain present across layers, superficial layers preferentially reflect top-down influences. Consequently, signals transmitted from S2 and posterior insula to other e.g. higher-order evaluative brain regions, likely already incorporate contextual and cognitive influences rather than representing purely sensory input. Sensory-discriminative cortices thus act as active nodes in cognitive pain modulation rather than passive relays.

At the same time, the similarity between bottom-up and top-down laminar profiles - both exhibiting inverted U-shaped patterns - suggests that cognitive modulation operates at multiple levels of the pain pathway. Because nociceptive stimulation was present in all conditions, feedforward input to cortex likely already carries information about cognitive state, consistent with evidence for modulation of nociceptive signals at subcortical stages, including the thalamus and spinal cord^18,33,34^.

This early modulation provides a parsimonious explanation for the shared features of the laminar profiles and motivates our focus on their relative differences rather than their absolute shapes^15,35^. Within this framework, cortical cognitive modulation reflects the combined influence of attenuated feedforward drive and altered cortico-cortical feedback. Our results provide evidence for the latter, while remaining compatible with parallel early suppression mechanisms. Together, these findings support a model in which cognitive modulation of pain emerges from interacting processes distributed across the neuraxis, with laminar-specific cortical dynamics shaping how these signals are integrated and propagated.

To address the inherent venous bias of GE-EPI^21^, we applied a factorial manipulation of nociceptive input and cognitive demand. Importantly, our inferences were based on relative differences between laminar profiles - “contrasts of contrasts” - rather than on their absolute shapes, thereby reducing sensitivity to vascular biases^15,35^. This distinction is critical, as sensitivity analyses indicate that we cannot fully exclude non-neuronal contributions to the observed inverted U-shaped laminar profiles (Supplementary Figures 8–9). Future work would benefit from complementary approaches that could enhance a more physiologically plausible separation of neural processing from vascular confounds, such as vascular space occupancy (VASO) imaging^36^ or laminar dynamic causal modeling^21,37^. In addition, although individual conditions elicited both positive and negative BOLD responses, their laminar dependencies were the same, consistent with previous observations in other sensory domains^38^.

Building on the present findings, several methodological and conceptual directions emerge for future layer-resolved investigations of pain. This approach could be used to disentangle individual differences in pain modulation mechanisms, such as the relative contribution of cortico-cortical feedback versus earlier, possibly cortico-spinal, modulation of nociceptive signals. Future work may benefit from refined experimental designs with illusory pain perception^39,40^ or targeted clinical populations with chronic or phantom limb pain to allow a cleaner dissociation between BU nociceptive drive and TD modulation. Future studies could integrate computational models of pain perception with laminar measurements^32,41,42^ to directly test the neural mechanisms proposed by predictive coding frameworks. Finally, our focus on individuals exhibiting robust behavioral analgesia reflects a deliberate emphasis on effective cognitive modulation but raises important questions about generalizability. Whether similar laminar mechanisms support other forms of pain modulation, such as nocebo hyperalgesia or placebo analgesia^2^, remains an open and promising avenue for future research. Together, these considerations highlight the potential of layer-specific fMRI to advance mechanistic understanding of pain and pave the way to individualized, circuit-informed approaches to pain modulation.

## Conclusion

Pain perception is a multifaceted process that integrates nociceptive, emotional, and cognitive components across multiple levels of the neuraxis. Here we show that high resolution fMRI enables investigation of layer-specific cortical processes underlying pain perception. Our study reveals distinct laminar profiles of BU and TD processes, indicating that sensory-discriminative regions integrate cognitive influences to shape the individual experience of pain. These findings highlight cortico-cortical feedback signalling as a mechanism underlying the cognitive modulation of pain.

## Methods

### Study design

In this preregistered study (https://osf.io/68je5), we investigated cortical laminar responses associated with cognitive (top-down effect, TD) and sensory modulation (bottom-up effect, BU) of pain. The experiment was conducted across three sessions on separate days (see Supplementary Fig. 1 and *Supplementary Methods—Study design*). During the first session, participants completed a psychophysical assessment to quantify individual behavioral analgesia induced by cognitive modulation. Only participants who met the preregistered criterion for behavioral analgesia were invited to the fMRI sessions (see *Behavioral paradigm*). The second session consisted of the acquisition of a high-resolution anatomical image (see *MRI acquisition*). In the third session, participants performed the experimental task during high-resolution fMRI data acquisition. In total, 34 participants completed the task-based fMRI session. Based on our preregistered inclusion criterion (behavioral analgesia must be present during the fMRI session), data from 30 participants were included in the final analysis. Sessions were separated by no more than 21 days (median = 11 days; range = 5–21 days).

### Participants

Healthy volunteers were recruited for this study; detailed inclusion and exclusion criteria are reported in the preregistration (https://osf.io/68je5). Based on an a priori sample-size estimation, a minimum of 30 participants was planned. In total, 89 individuals completed the initial psychophysical session (see *Study design*). According to the preregistered criteria (sufficient behavior distraction analgesia: change in the visual analoge scale (VAS) as a result of cognitive demand: ΔVAS_distraction_> 5, see *Behavioral paradigm*), 34 participants proceeded to the fMRI sessions. An additional exclusion criterion was preregistered based on behavior in the MRI session: only participants showing behavioral analgesia during scanning (ΔVAS_distraction_ > 0) were included in the analysis. The final sample therefore comprised data from 30 participants. To minimize expectancy effects related to pain modulation, the link between task performance and pain was not explicitly disclosed to participants.

### Behavioral paradigm

The experimental paradigm combined two forms of pain modulation: sensory (temperature; BU) and cognitive modulation (cognitive demand; TD), each with two levels, resulting in a 2 × 2 full-factorial design. The four experimental conditions were high demand/high temperature (HD/HT), low demand/high temperature (LD/HT), high demand/low temperature (HD/LT), and low demand/low temperature (LD/LT). Nociceptive heat stimulation was delivered to the left lower leg using a contact thermode (TSA2; ATS thermode; MEDOC, Israel; baseline temperature = 35 °C). Temperature levels were individually calibrated to elicit moderate and high pain (VAS40 and VAS70; see *Task and stimuli*). Cognitive modulation was induced using a modified n-back task (0-back: low demand; 2-back: high demand), as previously described^18^. The task was administered in a block design. Each block comprised an anticipation period, concurrent n-back task performance and heat stimulation, a rating period, and an inter-stimulus interval (Fig. 1A). Block duration was fixed at one of four values: 57, 60, 63, or 66 s. Each run consisted of 20 blocks, and participants completed three runs during the scanning session. To minimize sensitization and habituation, the thermode position was slightly adjusted between runs. Prior to the main task, participants completed an in-scanner practice session and an in-scanner calibration procedure (see *Supplementary Methods—MRI session*).

### Task and stimuli

Participants received detailed instructions regarding the task^18^, and before each block an instruction cue (0-back or 2-back) was provided about the upcoming task (*Supplementary Methods – Task and stimuli*).

During each task block, one of two temperature levels was applied, corresponding to moderate or high pain experience. After each task block (20 per run), participants rated the number of hits (range: 0–5) and the perceived pain intensity (VAS; anchors: 0 = not painful at all, 100 = extremely painful). Hit counts were reported using a forced-choice response between 0 and 5; however, the correct response always fell between 2 and 5. Participants were required to confirm their responses, with a maximum response window of 15 s. An additional inter-trial interval of 15, 18, 21, or 24 s followed each rating (Fig. 1A).

During the calibration procedure, participants received 10 heat stimuli at different temperature levels and rated the perceived intensity. The first highly salient stimulation was excluded and individual VAS40 and VAS70 temperatures were estimated using a regression-based approach ^43^. In a subsequent phase, each estimated temperature was delivered three times to verify that it reliably elicited the intended VAS levels. In cases of substantial mismatch, temperature values were adjusted and this phase was repeated. Stimulus presentation and task implementation were carried out using MATLAB (R2021b) and Psychtoolbox^44^.

### MRI acquisition

MRI data were acquired using a 7 T MRI system at the Ervin L. Hahn Institute (Essen, Germany) (Terra 7 T; Siemens Healthcare, Erlangen, Germany) equipped with a 1/32-channel transmit/receive radiofrequency (RF) head coil (NOVA Medical Inc.). For each participant, the vendor-provided 3rd order shimming was applied. Additional B1 maps were acquired using the preSAT-TFL technique ^45^ to calibrate transmitter amplitude, with the calibration region centered on the insula. Whole-brain T1-weighted anatomical images were acquired with an MP2RAGE sequence^46^ with fat navigators^47^ at 0.75-mm isotropic resolution. Parameter details can be found in Supplementary Table 1.

High-resolution functional task data were acquired with a 3D GRE-EPI sequence^48^. For each participant, the field of view (FOV) was angulated such that its inferior boundary was parallel to the most inferior aspect of the insula (*Supplementary Figure 10*). To compensate for imperfect slab excitation, the FOV was positioned slightly inferior to the insula. Fat suppression was achieved using a second-order water-excitation pulse (300 µs sub-pulse duration; bandwidth–time product = 23). Data acquisition was accelerated using CAIPIRINHA^49^. A GRAPPA autocalibration matrix of 96 x 48 lines was acquired in a separate EPI reference scan with a flip angle of 10°^50^. Parameter details can be found in Supplementary Table 1.

Single-coil images were combined using the vendor-provided adaptive combine algorithm, including virtual-adjust coil sensitivity phase correction, and coil-combined magnitude and phase data were saved for subsequent processing (see *MRI preprocessing*).

For distortion correction, each functional run was preceded by a sequence of inverted primary PE direction (∼1 min (13 volumes)), using otherwise identical parameters. Each scan began with three dummy volumes to allow magnetization to reach steady state; these volumes were discarded from all subsequent analyses.

### MR image processing

#### Preprocessing

Anatomical images were first reconstructed from raw k-space data using fat navigators (https://github.com/dgallichan/retroMoCoBox) for motion correction. In the next step, brain extraction (https://github.com/srikash/presurfer) was performed and manually inspected to correct erroneous brain masks. White-matter segmentation was performed with Freesurfer (https://surfer.nmr.mgh.harvard.edu/) and used for component-based noise correction (CompCor)^51^ within the general linear model (GLM) framework (see *General linear model on imaging data)*. In case of an erroneous brain extraction, masks were manually corrected.

The masked and motion-corrected images were then processed with the Freesurfer recon-all pipeline^52^ to reconstruct white-matter and pial surfaces. For each individual, the white matter surface was manually inspected in the primary and secondary somatosensory cortices and the insula (*Supplementary Figure 3*), defining our region of interests (ROIs). In cases of surface misdelineation, which occurring more frequently in the insula, surfaces were manually corrected following our previous work^53^. The Freesurfer pipeline was re-run on the corrected images and this procedure was repeated iteratively until satisfactory surface reconstruction was achieved. Laminar analyses were performed in individual anatomical space (see *Laminar sampling*). Additionally, the inverse transformation of a nonlinear registration from individual anatomy to a common template (tpl-MNI152NLin2009cAsym_res-01; https://github.com/templateflow) was computed and later used for generating group level activation maps.

For the functional scans, partial-Fourier reconstruction was performed offline using the complex data and the projection onto convex sets (POCS) algorithm^54,55^ to preserve spatial resolution (see the preregistration for details). The POCS-reconstructed task fMRI data were preprocessed, including motion correction, distortion correction, and registration to the anatomical image. ANTS^56^, FSL^57^ and SPM12^58^ were used within our in-house processing pipeline (https://github.com/kincsesbalint/LaminarfMRI_inPainPerception) to calculate individual spatial transformations. Resampling with Lanczos windowed sinc interpolation was applied in a single step (ANTS). First level GLM parameters were estimated in individual anatomical space (*General linear model on imaging data*).

#### General linear model on imaging data

The first level results were estimated with a GLM as implemented in SPM. A high-pass filter was applied (cut-off: 180 s) and the prewhitening was performed using an AR(1) model. The design matrix contained seven regressors per run, including anticipation periods of the two task types (see *Behavioral paradigm*), the four experimental conditions (high demand/high temperature (HD/HT), low demand/high temperature (LD/HT), high demand/low temperature (HD/LT), low demand/low temperature (LD/LT)), and the rating period. Six motion parameters, and the first five components of the white matter anatomical CompCor were included as additional nuisance regressors and all regressors were convolved with the canonical hemodynamic response function (HRF).

The parameters of the main effect of temperature and cognition were estimated by using the robust weighted least squares toolbox (rWLS)^59^ for the unsmoothed, laminar level (averaged across vertex pairs, see below) timeseries data, with fixed effects estimation across runs. As an initial step in the functional ROI definition, the restricted Maximum Likelihood estimation was used on the smoothed (4 mm kernel) high-resolution data in individual space (see *Functional ROI definition*). That is the main effect of temperature was defined as the difference of high temperature and low temperature (HT - LT) averaged across distraction levels, and the main effect of distraction was calculated as the difference of low demand/weak distraction and high demand/strong distraction (LD - HD) averaged across temperature levels. These contrast estimates (main effect of temperature and distraction) and corresponding T-statistics of the main effects were used in later steps of the analysis.

Additionally, a second level GLM was applied to estimate group-level activation (*Supplementary Figure 7*). These activations (thresholded at t > 1, uncorrected) was used to define functional volumetric ROIs (see *Functional ROI definition*). Timeseries extraction was performed between the WM/CSF vertex pairs within these ROIs (see *Laminar sampling*).

#### Functional ROI definition

To identify individual functional ROIs for further analysis, a combined strategy incorporating anatomical information, group level activation maps, and individual level activation maps was used (https://osf.io/68je5). In detail, the group-level activation maps of the main effects (temperature and distraction) were used in a conjunction analysis^60^, to restrict subsequent analyses to cortical regions that responded to nociceptive input and also showed modulation by distraction (*Supplementary Figure 7*). This map was first intersected with anatomical regions (S1, S2, pIns; *Supplementary Figure 3*) based on Schaefer atlas (200 parcels), and subsequently dilated and binarized. The resulting functional ROI was then transformed back to the individual space. Left and right hemispheres were kept separately during the analysis pipeline because of the known asymmetry of pain elicited brain signal^61^. The individual functional ROI volumes were then projected to the WM surface (Workbench Command) resulting in a set of vertices serving as functionally defined masks in surface space with their corresponding vertex pairs in the pial surface. All subsequent analyses were restricted to the subset of vertices comprising this WM surface mask (Figure 1C, see *Laminar sampling*).

#### Laminar sampling

We describe below a method for selecting functionally relevant surface elements (i.e.: vertex pairs) within the previously defined surface ROIs. Importantly, the definition of the surface ROIs using this method was orthogonal to the main research question, which investigates the laminar differences of the effects. The individual surface ROIs consisted of sets of vertex pairs connecting the WM and pial surfaces. Along these vertex pairs, the preprocessed fMRI timeseries data were sampled equidistantly (SPM, https://github.com/kincsesbalint/LaminarfMRI_inPainPerception) at 20 data points using a third-order polynomial interpolation. Note that these datapoints define the “layers”; they do not correspond to histologically defined cortical layers, as the current spatial resolution of laminar fMRI is insufficient to resolve individual cytoarchitectonic layers.

To select vertices for averaging, a vertex-wise conjunction analysis^60^ was performed using the statistics of the parameter estimates for both BU (temperature) and TD (distraction). To reduce the superficial bias of the BOLD signal^14,62,63^, the timeseries from the deeper cortical depths (i.e., from WM surface to 75 % of the cortical depth toward the pial surface) were averaged and the same first-level GLM as in voxel space was fitted. The top 200 vertex pairs were selected, based on their positive vertex-wise t-values derived from the conjunction analysis (uncorrected, see also *Sensitivity analysis* for further selection criteria).

In the next step, timeseries signals were averaged across vertices from the selected 200 “columns”. The main effects of temperature and distraction were then estimated separately for each layer (20 layers, see *First level estimation*). The resulting individual contrast estimates were subsequently used in a statistical analysis (20 values across the cortical depth x 2 processes per individual).

### Sensitivity analysis

To investigate the robustness of our findings, we conducted additional sensitivity analyses. Specifically, within the functionally defined ROIs (based on the group-level activation) the selection of vertices at the individual level was investigated. In one approach, we applied a more liberate threshold for the selection of vertex pairs (“columns”), including all vertices with positive vertex wise t-values (t > 0). That is, all vertices showing effects of temperature and distraction in the expected direction (higher activation with higher nociceptive input and higher activation under lower distraction) were included. Importantly, this approach resulted in a larger number of included vertices per participant, which reduces potential samping bias across the cortical depth^64^.

The second approach focused on vertex selection based on smoothed individual level data. In this case, activation maps derived from smoothed individual images were thresholded at t > 1 for the conjunction of temperature and distraction effects (see above). The resulting ROI was projected onto the cortical surface, and the averaged timeseries signal across these vertices was calculated and introduced into a GLM. Results from these sensitivity analyses are reported in the *Supplementary results* and *Supplementary Figure 8-9*.

### Statistical analysis

All statistical modeling was performed in R^65^ and is available on the project’s GitHub repository (https://github.com/kincsesbalint/LaminarfMRI_inPainPerception). Behavioral and imaging data were modeled using a random-intercept linear mixed-effects model^66,67^. Pain-intensity ratings served as the response variable, temperature and cognitive demand were included as fixed factors, and participants were modeled as a random effect. The significance of model parameters^67^ and their confidence intervals were estimated using the parameters package^68^. Equal-tailed uncertainty intervals and two-tailed p-values were computed using a Wald t-distribution approximation. Individual behavioral distraction and temperature effects were calculated by averaging observations within each condition (five observations per run × 3 runs = 15) and computing the effect of distraction and temperature by contrasting the corresponding conditions (Equation 1,2).

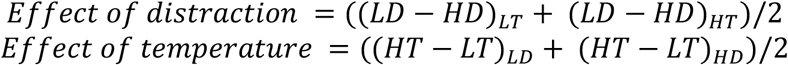

*Equation 1 and 2 Calculation of cognitive demand/distraction and temperature effects. The VAS ratings were used in each condition, and their arithmetic differences was derived to quantify the effects of cognitive demand and temperature. LD = low demand, HD = high demand, LT = low temperature, HT = high temperature. The same equations were used for the imaging data, that is contrast estimates derived from the ß estimates*.

In the imaging data, the exploratory nature of the project motivated a more liberate statistical approach, therefore, each ROI was handled separately, and linear mixed-effects models were fitted independently^66,67^. Brain activity (contrast estimates) served as the response variable, and ‘process’ (levels: bottom-up/effect of temperature, top-down/effect of distraction), ‘layer’ (handled as a continuous variable), and their interaction were included as explanatory variables. A quadratic term of layers was also included (polynomial model), to capture the inverted U-shaped profile observed in the data. The inclusion of the quadratic term significantly improved model fit, as confirmed by model comparison with ANOVA (see *Supplementary methods - statistical modeling*, https://github.com/kincsesbalint/LaminarfMRI_inPainPerception/, *Supplementary results – Model fitting*). Because a quadratic term for layer was included, the process × layer interaction comprised both linear and quadratic components (see *Supplementary methods - statistical modeling)*.

Fixed effects were evaluated with Type III F-tests with Satterthwaite’s degrees-of-freedom approximation. Following the analysis strategy of Lawrence et al.^15^, a significant ‘process x layer’ interaction effect was interpreted as evidence for differential involvement of cortical layers in top down and bottom up pain processing.

In addition, we fitted a random intercept model treating ‘layer’ as a categorical variable, using the layer closest to the white matter (layer 1) as reference level, to identify which specific layers drove the interaction. Because this specification estimates a larger number of parameters than the polynomial model, it is statistically less efficient (i.e., lower power) for detecting differences. We therefore used the polynomial model for primary inference and the categorical model for descriptive localization, with correction for multiple comparison. Post-hoc comparison of fixed-effect coefficients for processes were tested using two-sided t-tests with Satterthwaite’s degrees-of-freedom approximation (Type III).

To derive individual differences in laminar profiles between BU and TD processes, we additionally repeated model fitting with layer as a continuous variable and included random slopes for the interaction term. Because individual laminar profile shapes are defined by all the coefficients of the second-order polynomial, an area-under-the-curve (AUC) metric was computed to capture and quantify overall laminar profile for each process and each individual. We calculated the difference of the AUC_TD_ and AUC_BU_, which we refer to as the *individual relative agranular involvement*. Higher ΔAUC values (interpreted already the fixed effect coefficients from the model) indicate greater relative agranular involvement during TD process. To investigate the association between different laminar signaling and behavior, the individual ΔAUC measure (*individual relative agranular involvement*) was subsequently entered into a correlation analysis with behavior analgesia (two-sided Spearman test).

Statistical significance was set at p = 0.05. Given the exploratory nature of the study and the need to balance Type I and Type II errors, we did not apply corrections for multiple comparisons across regions; however, false discovery rate (FDR) correction was applied within each “localization” model (i.e., identifying significant levels of the layer variable in the categorical models; see above).

## Data availability

The used codes and data are available in the project OSF page (https://osf.io/ht3dw/) and github page (https://github.com/kincsesbalint/LaminarfMRI_inPainPerception/).

## Acknowledgements

The work is funded by the Deutsche Forschungsgemeinschaft (DFG, German Research Foundation) – Project-ID 422744262 – TRR 289 (“Treatment Expectation”) (Gefördert durch die Deutsche Forschungsgemeinschaft (DFG) – Projektnummer 422744262 – TRR 289), Project-ID 316803389 – TRR 1280 (“Extinction Learning”). The MAGNETOM Terra used in the study was funded by the Deutsche Forschungsgemeinschaft (DFG, German Research Foundation), grant number 432647511. KW was supported by the National Institute for Health and Care Research (NIHR) Oxford Health Biomedical Research Centre (OH-BRC). The views expressed are those of the authors and do not necessarily reflect those of the NIHR or the Department of Health and Social Care. The authors are thankful to Prof. David Norris for the discussion of methodological considerations and the results.

## Author contributions

B.K., V.P., P.K. and U.B. conceived the study. The study was performed by B.K., V.P., K.P. B.K. and V.P. conducted the analyses with the supervision of T.S. and P.K.; B.K.,V.P., K.W., and U.B. wrote the manuscript. B.K. V.P., T.S. K.W., P.K., and U.B. contributed to the interpretation, as well as manuscript revision. The whole study was supervised by P.K. and U.B.

