## Supplementary for "Layer-specific cortical processing dissociates sensory and cognitive influences on pain"

### Supplementary methods

#### Study design

This study took place on three separate days (Supplementary Figure 1). On the first day (psychophysical session), participants performed the cognitive task in conjunction with a painful stimulus. Before the main task, participants completed a practice task and a calibration procedure (see *Psychophysics session*), and completed additional psychological questionnaires (data has not been used in this analysis).

The MRI sessions were conducted on two separate days, during which participants completed a resting-state fMRI session and a task-based fMRI session. During the resting-state fMRI session, a high-resolution anatomical image and a resting-state fMRI scan were acquired (see *MRI acquisition* and *Supplementary Table 1*). In addition, quantitative sensory testing was performed outside the scanner following the MRI session. Only the anatomical data acquired on this day were used in the current analyses.

During the task-fMRI session, participants performed the experimental task during high-resolution functional MRI acquisition. As in the psychophysical session, a practice task and a calibration procedure preceded the main task.

#### Participants

In total, 89 participants were recruited, of whom 36 were invited to participate in the MRI sessions. One participant withdrew from the study before the MRI sessions, and another completed only the resting-state fMRI session; this resulted in a final sample of 34 participants with complete imaging and behavioral data. Participants were informed that the primary aim of the study was to investigate the neural mechanisms underlying task performance and their relation to psychological states and traits, which were assessed using questionnaires (data has not been used in this analysis). Participants were additionally informed that painful stimulation was included to ensure comparability with previous studies, rather than as the central focus of the experiment, to avoid contextual effects of cognitive modulation.

#### Task and stimuli

On day 1, prior to the main paradigm, participants practiced the n-back task (practice phase) until they achieved successful performance in at least three of the last five blocks (defined as a correct report of the number of hits). Because participants had already reached adequate task performance during the first session, practice on day 3 always consisted of six blocks in the scanner.

The paradigm followed a block design structure, with each block comprising an anticipation phase, the n-back task combined with painful stimulation, a rating period, and an interstimulus interval (Figure 1A). Participants were instructed to count the number of hits in each block. To simplify instructions, the tasks were referred to as the "Search" and "Remember" tasks for the 0-back and 2-back conditions, respectively. Each n-back block began with an instruction cue indicating either "search" (0-back) or "remember" (2-back). In the 0-back condition, an additional letter was presented as the target, and participants counted its occurrences. In the 2-back condition, participants monitored whether the current letter matched the letter presented two trials earlier and counted such matches as hits. Each letter was displayed for 0.75 s, with an inter stimulus interval of 0.75 s. Each block contained 16 letters, resulting in a 24 s task duration. Letters ranged from A to E and were presented in a pseudorandom sequence.

Following the practice phase, heat pain thresholds were determined, and a calibration procedure was conducted. During calibration, participants received 19 stimuli at different temperatures on day 1, or 10 stimuli during scanning on day 3. The first rating was excluded, and a regression based approach was used to estimate temperatures corresponding to VAS40 and VAS70. In a subsequent post calibration phase, these two temperatures were each repeated three times to verify the accuracy of the estimates.

Task conditions were presented in a pseudorandomized order, using ten predefined sequences comprising four conditions: high temperature/low demand (0-back), high temperature/high demand (2-back), low temperature/low demand (0-back), and low temperature/high demand (2-back). Each run contained five blocks per condition, with the constraints that high-pain conditions were repeated no more than twice consecutively and high-demand conditions no more than four times consecutively. For each participant, five runs (two on day 1 and three on day 3) were randomly selected from these sequences. Pilot data showed no sequence effects on pain ratings.

#### Task fMRI session

The practice of the task in the scanner, conducted without concurrent heat pain stimulation, consisted of six blocks and was intended to facilitate adaptation to the scanner environment and familiarization with the MRI compatible response pad.

During the heat calibration phase, individual stimulus temperatures were adjusted to levels perceived as VAS40 and VAS70 on a visual analogue scale (VAS) ranging from 0 to 100 (anchors: 0, not painful at all; 100, highest imaginable pain). The resulting temperatures were used throughout the main task.

Additional functional scans were acquired during the practice and calibration phases of the task fMRI session. In our preregistered approach for defining functional regions of interest (ROIs), these scans were initially intended for use as functional localizers. However, suboptimal contrasts and limited scan duration reduced their reliability, limiting their suitability for ROI definition. We therefore adopted an alternative, preregistered approach to identify individual functional ROIs (see *Methods*)

Each functional scan was preceded by an acquisition with reversed phase encoding direction, which was used to estimate the field map for distortion correction and image registration.

Regarding the main task, the same experimental paradigm was used in both the psychophysical and MRI sessions (see *Methods—Task and stimuli*).

#### Statistical modelling

A linear mixed-effects model was fitted with fixed effects of process (two levels: cognitive demand and nociception), layer (modeled either as a continuous variable or as a categorical variable with 20 levels), and their interaction, and with a random intercept for subject. Using the Wilkinson–Rogers notation in an lme4 framework, the following models were specified:

Random-intercept model:

$$contrast\ estimate \sim process * (\,layer\ +\ I(layer^2)) + (1|subj)$$

Random-slope model:

$$contrast\ estimate \sim process\ *\ (\,layer\ +\ I(layer^2))\ +\ (1 +\ process{:}(\,layer\ +\ I(layer^2))|\ subj)$$

To localize the cortical layers driving the interaction, an additional model was fitted in which layer was treated as a categorical variable, with the deepest layer serving as the reference level:

$$contrast\ estimate \sim process * layer\ + (1|subj)$$

This categorical model has reduced statistical power compared with the random-intercept (continuous) model. The analysis code is publicly available on the project's GitHub repository (https://github.com/kincsesbalint/LaminarfMRI_inPainPerception/).

### Supplementary results

#### Sensitivity analysis

We performed additional sensitivity analyses to assess the robustness of results to individual-level selection of active regions, specifically the definition of active vertices. In one approach, all vertices showing a positive effect for both BU and TD processes were included in the analysis. This increased the number of included vertices. This analysis revealed a similar laminar activation pattern (Supplementary Fig. 8, middle row), with significant interaction terms for both the linear and quadratic components, and the association with behavioral analgesia remained observable in S2 and posterior insula (pIns) (Supplementary Fig. 8, bottom row).

In a second, more liberal inclusion approach, individually smoothed images thresholded at $t > 1$ on the individual conjunction maps (see *Methods*) were used to define regions of interest for the laminar pattern-difference analysis. Under this criterion, individual laminar profiles exhibited a monotonic increase toward superficial layers, while a robust difference between the two processes was preserved in superficial layers (Supplementary Fig. 9, middle row). Specifically, TD processing was associated with stronger superficial-layer activation, as reflected by significant interaction terms in four of six regions of interest. The correlation between laminar pattern differences and behavioral analgesia remained significant in S2 (Supplementary Fig. 9, bottom row). Note that vertex selection under this approach may be biased by signals originating outside the functional ROI. In particular, pial veins lie outside the gray-matter ribbon, and their contribution can be amplified by spatial smoothing. In addition, signals from the gray matter on the opposite bank of a sulcus, may also influence vertex selection.

#### Model fitting

The inclusion of a quadratic term in the linear mixed-effects model was motivated by visual inspection of the laminar response profiles and formally evaluated through statistical model comparison. Across all regions of interest, model fit improved significantly with the addition of the quadratic term, supporting its inclusion in subsequent analyses.

```
[1] "The S1 region and left side"
Models:
model_linear: beta ~ process * layeridx_scaled + (1 | subjid)
model_quadratic: beta ~ process * (layeridx_scaled + I(layeridx_scaled^2)) + (1 |
subjid)
                npar    AIC    BIC  logLik -2*log(L)  Chisq Df Pr(>Chisq)
model_linear       6 8334.9 8365.4 -4161.4    8322.9
model_quadratic    8 8052.2 8093.0 -4018.1    8036.2 286.64  2  < 2.2e-16 ***
---
Signif. codes:  0 ‘***’ 0.001 ‘**’ 0.01 ‘*’ 0.05 ‘.’ 0.1 ‘ ’ 1
--------------------------------------------------
[1] "The S1 region and right side"
Models:
model_linear: beta ~ process * layeridx_scaled + (1 | subjid)
model_quadratic: beta ~ process * (layeridx_scaled + I(layeridx_scaled^2)) + (1 |
subjid)
                npar    AIC    BIC  logLik -2*log(L)  Chisq Df Pr(>Chisq)
model_linear       6 9134.4 9164.9 -4561.2    9122.4
model_quadratic    8 8783.9 8824.6 -4383.9    8767.9 354.51  2  < 2.2e-16 ***
---
Signif. codes:  0 ‘***’ 0.001 ‘**’ 0.01 ‘*’ 0.05 ‘.’ 0.1 ‘ ’ 1
--------------------------------------------------

[1] "The S2 region and left side"
refitting model(s) with ML (instead of REML)Data: moddata
Models:
model_linear: beta ~ process * layeridx_scaled + (1 | subjid)
model_quadratic: beta ~ process * (layeridx_scaled + I(layeridx_scaled^2)) + (1 |
subjid)
                npar    AIC    BIC  logLik -2*log(L)  Chisq Df Pr(>Chisq)
model_linear       6 8571.6 8602.1 -4279.8    8559.6
model_quadratic    8 7953.5 7994.2 -3968.7    7937.5 622.07  2  < 2.2e-16 ***
---
Signif. codes:  0 ‘***’ 0.001 ‘**’ 0.01 ‘*’ 0.05 ‘.’ 0.1 ‘ ’ 1
--------------------------------------------------
[1] "The S2 region and right side"
Models:
model_linear: beta ~ process * layeridx_scaled + (1 | subjid)
model_quadratic: beta ~ process * (layeridx_scaled + I(layeridx_scaled^2)) + (1 |
subjid)
                npar    AIC    BIC  logLik -2*log(L)  Chisq Df Pr(>Chisq)
model_linear       6 9049.6 9080.1 -4518.8    9037.6
model_quadratic    8 8389.5 8430.2 -4186.8    8373.5 664.06  2  < 2.2e-16 ***
---
Signif. codes:  0 ‘***’ 0.001 ‘**’ 0.01 ‘*’ 0.05 ‘.’ 0.1 ‘ ’ 1
--------------------------------------------------
```

```
[1] "The pIns region and left side"
refitting model(s) with ML (instead of REML)Data: moddata
Models:
model_linear: beta ~ process * layeridx_scaled + (1 | subjid)
model_quadratic: beta ~ process * (layeridx_scaled + I(layeridx_scaled^2)) + (1 |
subjid)
                npar    AIC    BIC  logLik -2*log(L) Chisq Df Pr(>Chisq)
model_linear       6 8242.6 8273.1 -4115.3    8230.6
model_quadratic    8 7460.7 7501.4 -3722.3    7444.7 785.9  2  < 2.2e-16 ***
---
Signif. codes:  0 ‘***’ 0.001 ‘**’ 0.01 ‘*’ 0.05 ‘.’ 0.1 ‘ ’ 1
--------------------------------------------------
[1] "The pIns region and right side"
Models:
model_linear: beta ~ process * layeridx_scaled + (1 | subjid)
model_quadratic: beta ~ process * (layeridx_scaled + I(layeridx_scaled^2)) + (1 |
subjid)
                npar    AIC    BIC  logLik -2*log(L)  Chisq Df Pr(>Chisq)
model_linear       6 8468.4 8498.9 -4228.2    8456.4
model_quadratic    8 7775.1 7815.8 -3879.6    7759.1 697.26  2  < 2.2e-16 ***
---
Signif. codes:  0 ‘***’ 0.001 ‘**’ 0.01 ‘*’ 0.05 ‘.’ 0.1 ‘ ’ 1
```

### Supplementary Figures

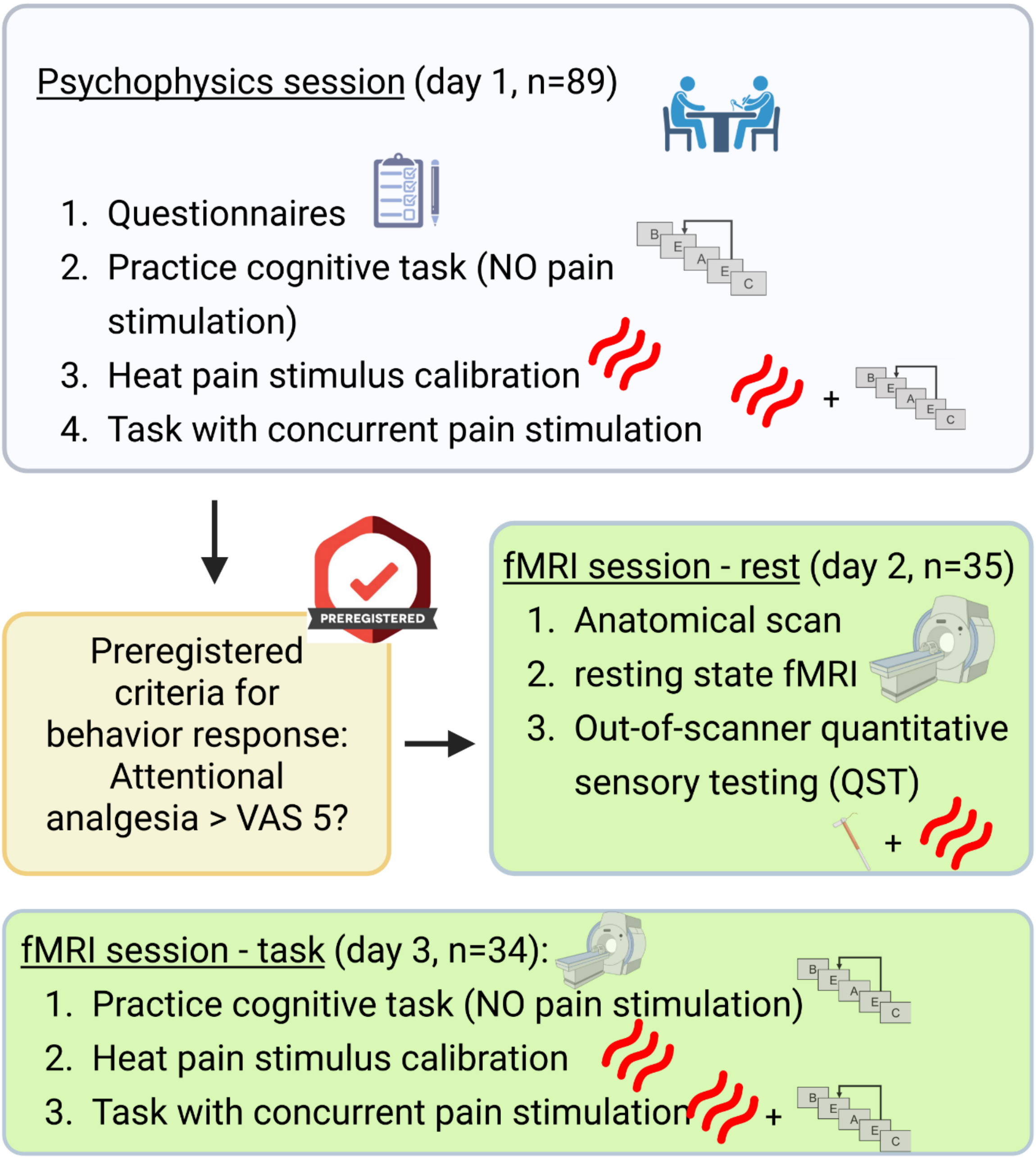


*Supplementary Figure 1 Overview of the study design across three experimental days.*
*The study was conducted over three separate days. On day 1 (psychophysical session), participants completed battery of questionnaires followed by task practice, heat-pain calibration, and the main behavioral paradigm outside the scanner. On day 2 (resting-state MRI session), high-resolution anatomical images and resting-state fMRI data were acquired; quantitative sensory testing was performed after scanning, although only anatomical data from this session were used in the present analyses. On day 3 (task-fMRI session), participants performed the same experimental paradigm during high-resolution fMRI acquisition, preceded by task practice and heat-pain calibration.*

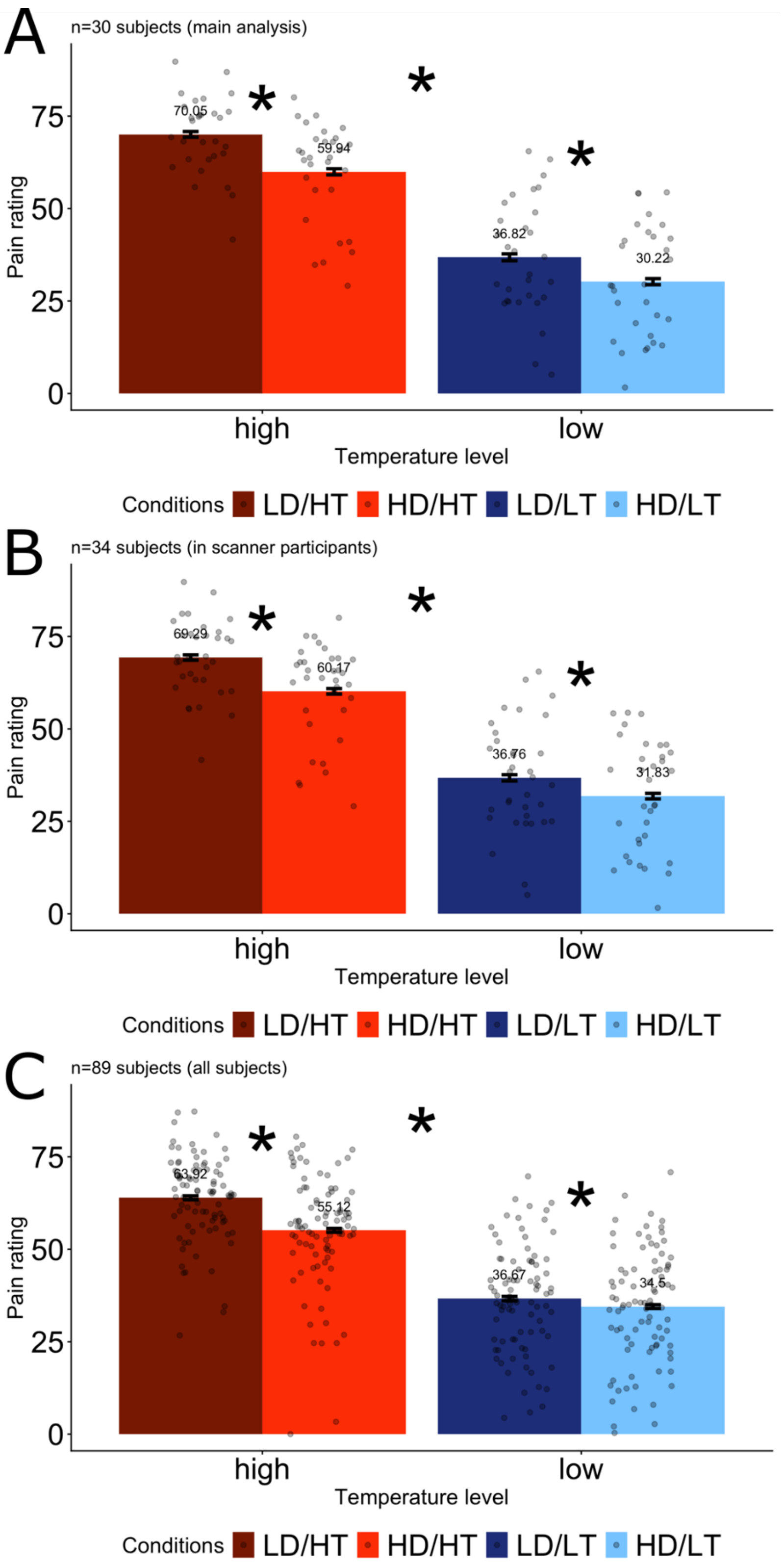


*Supplementary Figure 2 Group-level pain ratings across cohorts and experimental sessions. Bar plots show mean pain ratings for all experimental conditions at the group level. Panel A displays results for the cohort selected according to the preregistered criterion (attentional analgesia > 0, n=30), panel B shows data from the MRI session from all participants (n=34), and panel C shows data from the psychophysical session (n=89). Across all cohorts, a significant main effect of temperature, a significant main effect of task demand, and a significant temperature × demand interaction were observed at the group level. LD and HD denote low and high demand, respectively, and LT and HT denote low and high temperature.*

A) The used volumetric region of interests

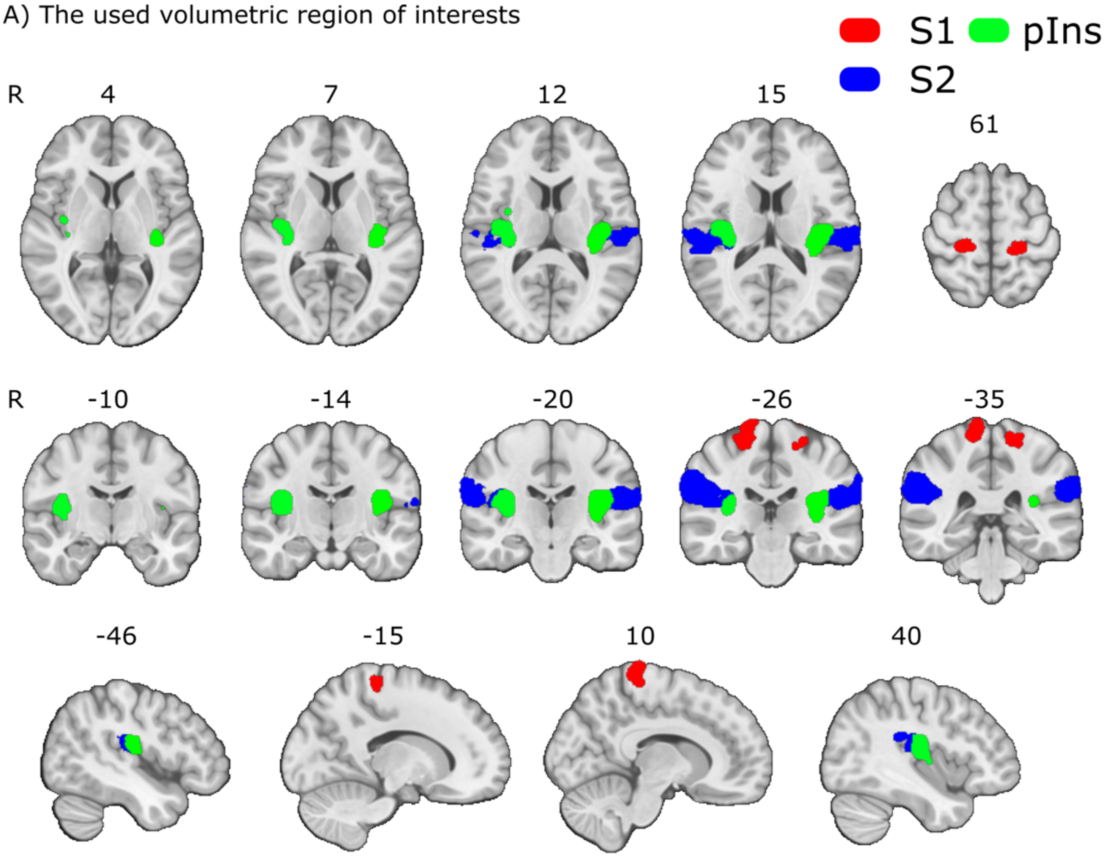


B) Brain activation (ß/con) values independent of layers in each ROI

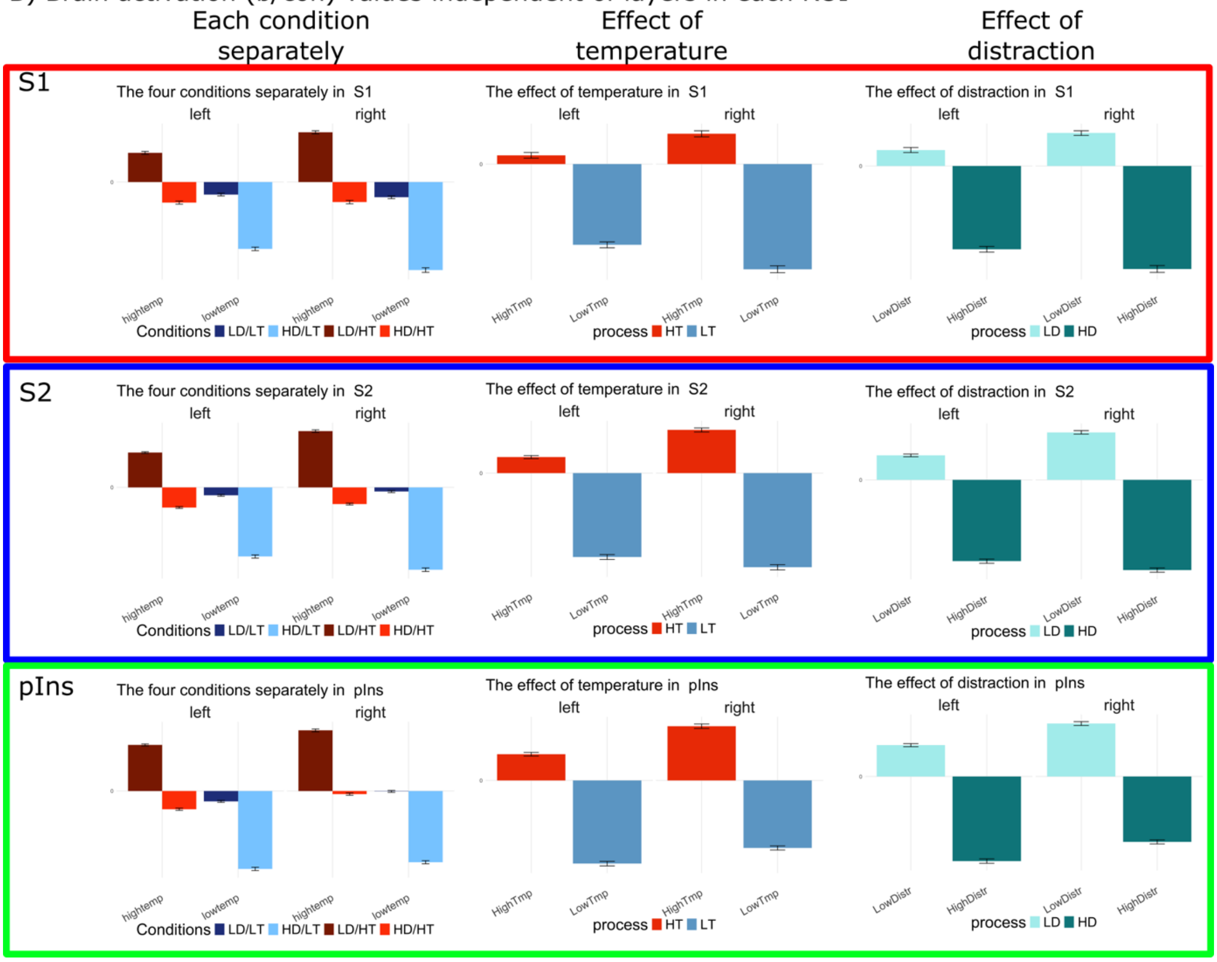

*Supplementary Figure 3 Regions of interest used in the analyses and corresponding group-level activation.*
*A) Anatomical regions of interest (ROIs) are shown in MNI space. The MNI coordinates of the center of gravity for each ROI are as follows (X, Y, Z): Left S1: −20.63, −31.56, 64.57; Right S1: 16.33, −30.70, 70.30; Left S2: −59.05, −26.71, 21.62; Right S2: 56.46, −27.92, 25.32; Left posterior insula: −37.70, −21.34, 14.72; Right posterior insula: 36.95, −17.60, 14.71.*
*B) Bar plots depict mean activation averaged across cortical layers for each experimental condition. Separate bar plots additionally illustrate the main effects of temperature and task demand. LD and HD denote low and high demand, respectively, and LT and HT denote low and high temperature.*

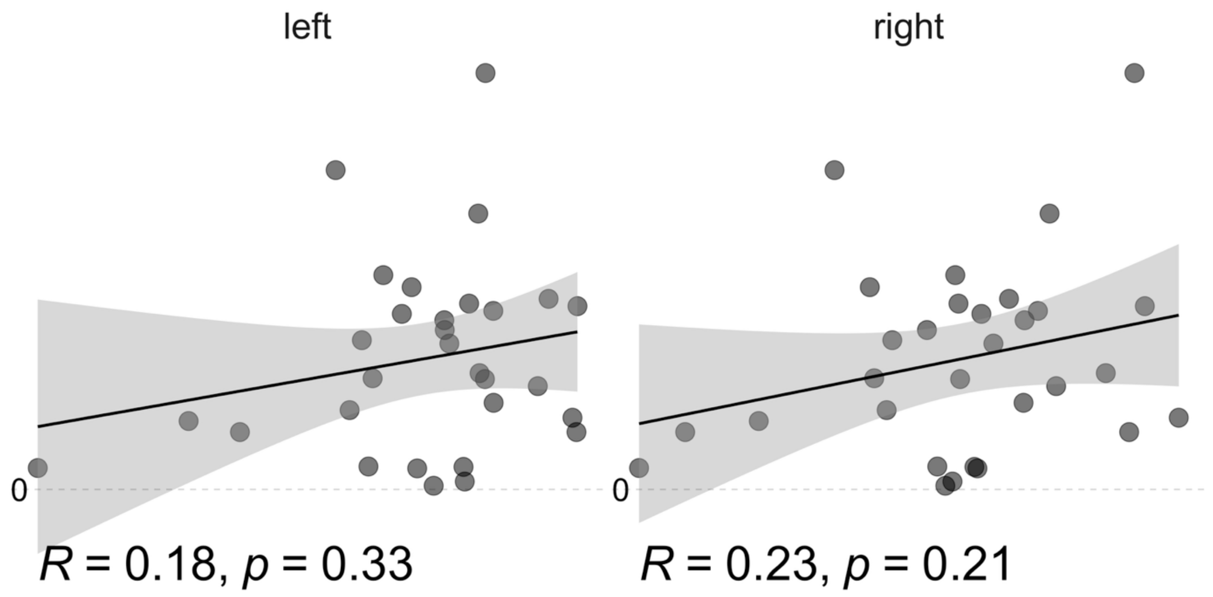


*Supplementary Figure 4 Relationship between relative agranular involvement in S1 and attentional analgesia. Scatter plot illustrating the correlation between the relative agranular involvement in primary somatosensory cortex (S1) (see Fig. 2 in the main manuscript for definition of the metric) and individual differences in attentional analgesia, quantified from behavioral pain ratings. Each data point represents one participant.*

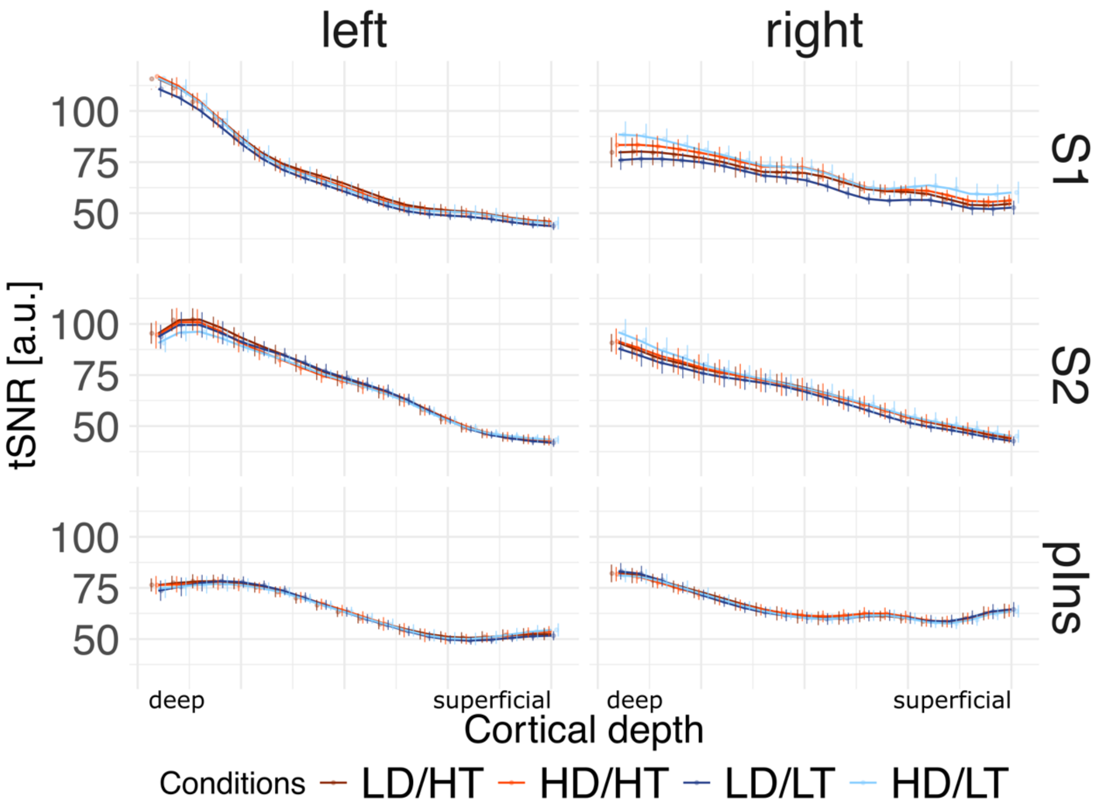


*Supplementary Figure 5 Temporal signal-to-noise ratio across cortical layers and conditions. Group-averaged temporal signal-to-noise ratio (tSNR) is shown for each experimental condition and region of interest (ROI), plotted separately for the left and right hemispheres. Error bars indicate the standard deviation. LD and HD denote low and high demand, respectively, and LT and HT denote low and high temperature. Across ROIs, tSNR showed decrease from deeper to more superficial layers (1: WM/GM boundary; 20: pial surface). No systematic differences in tSNR were observed between experimental conditions, although substantial interindividual variability was present.*

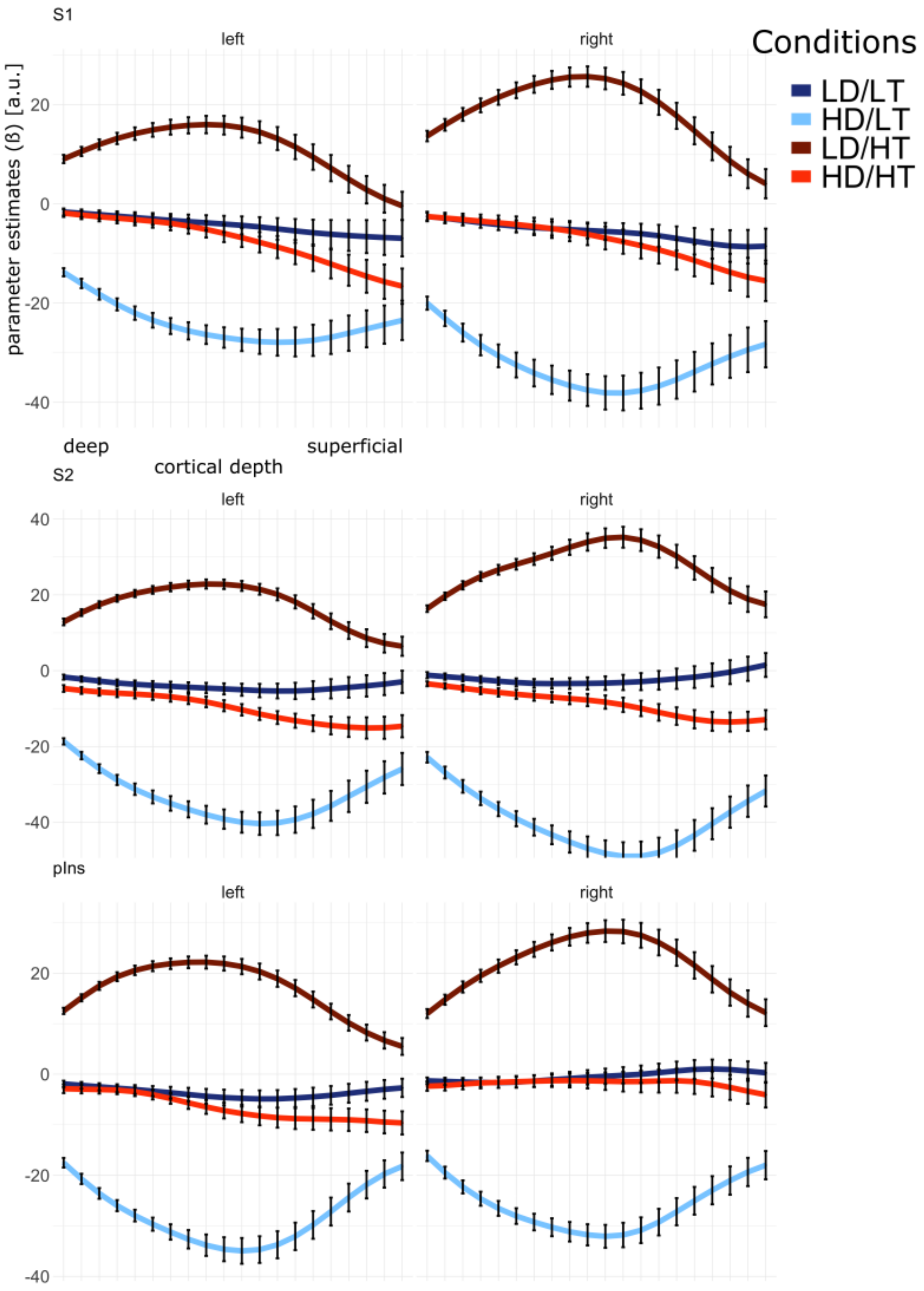


*Supplementary Figure 6 Layer-dependent parameter estimates across conditions and regions of interest. Condition-specific parameter estimates are plotted as a function of cortical layer for each region of interest (ROI). Red curves indicate high-temperature conditions (HD/HT and LD/HT), whereas blue curves indicate low-temperature conditions (HD/LT and LD/LT). Dark shades denote low-demand conditions, and light shades denote high-demand conditions. Points indicate the mean, and error bars represent the standard error of the mean.*

Group level activation maps:

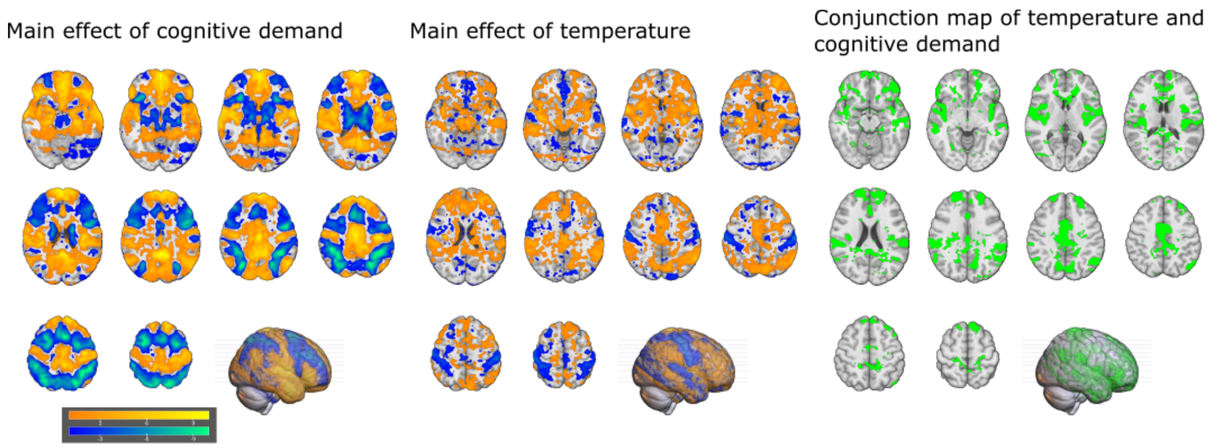


*Supplementary Figure 7 Main effects of task demand, temperature, and their uncorrected conjunction map. Statistical parametric maps illustrating the main effect of task demand (left), the main effect of temperature (middle), and their uncorrected conjunction (right). All maps are thresholded at t > 1. For the main effect of demand, yellow regions indicate relatively greater activation during low-demand compared with high-demand conditions (contrast: low demand – high demand > 0), whereas blue regions indicate relatively greater activation during high-demand compared with low-demand conditions (contrast: high demand – low demand > 0). For the main effect of temperature, yellow regions indicate increased activation for high compared with low temperature (contrast: high temperature – low temperature > 0), whereas blue regions indicate the opposite effect (contrast: low temperature – high temperature > 0). For the conjunction analysis, uncorrected t-maps for demand and temperature were combined using a conjunction approach (see Methods). Green regions indicate voxels showing both cognitive demand related and nociceptive effects. These regions were intersected with anatomical regions of interest (Supplementary Fig. 3) and subsequently used to define functional ROIs in individual subject space.*
*Axial slices are shown at z = −15, −6, 6, 15, 22, 30, 38, 45, 52, and 59 in MNI space. The original statistical maps are available on the project's OSF repository.*

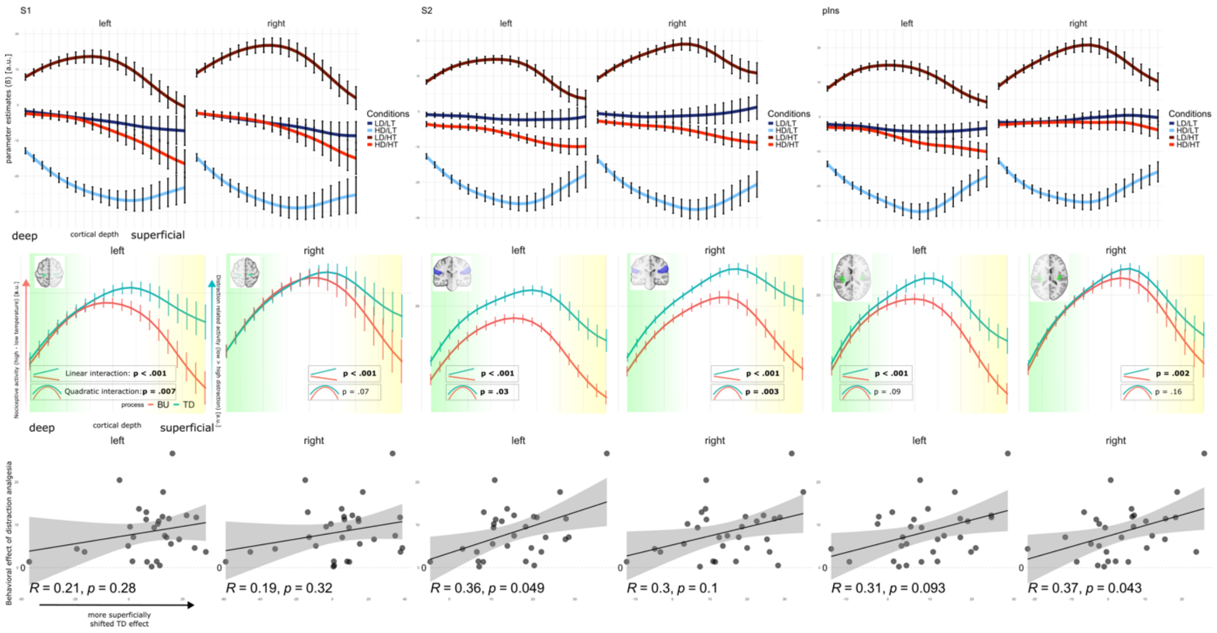


*Supplementary Figure 8 Sensitivity analysis of vertex-selection criteria.*
*Results obtained by including all positive vertices identified in the conjunction-based, vertex-wise analysis are shown (see Methods). A similar laminar response pattern was observed across all regions and hemispheres as in the main analysis (Figure 2), including the main-effect laminar distributions (second row). In addition, the laminar pattern difference (defined as relative agranular involvement; see Methods) was associated with the magnitude of behavioral analgesia in left S2 and right posterior insula, whereas comparable effects were observed in the contralateral regions. All reported P values are from two-sided statistical tests.*

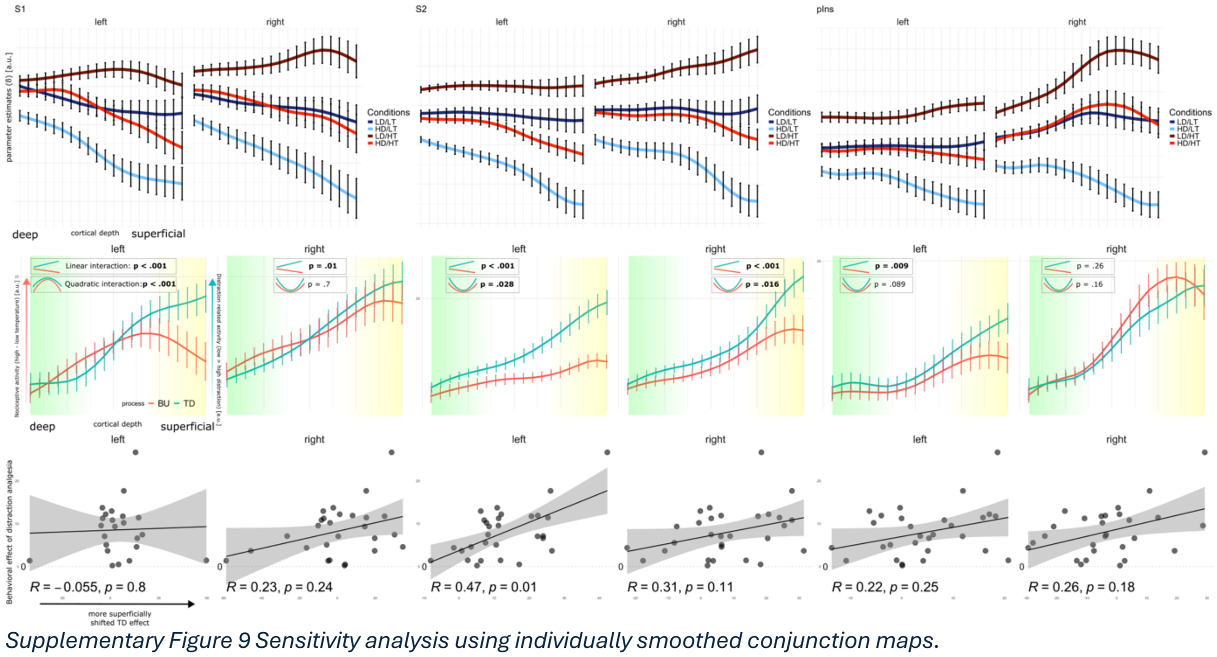


*Supplementary Figure 9 Sensitivity analysis using individually smoothed conjunction maps.*
*Conjunction maps were generated from individually smoothed images and thresholded at t > 1 (see Methods). Compared with the main analysis, individual process-related laminar profiles showed a shift toward a more linear increase across superficial layers. However, formal model-comparison analyses continued to favor the inclusion of a quadratic term in most regions (4 of 6 regions). Differences between the two processes remained most evident in left S1, S2, and left posterior Ins. In left S2, the significant correlation between laminar pattern differences and behavioral attentional analgesia was preserved.*

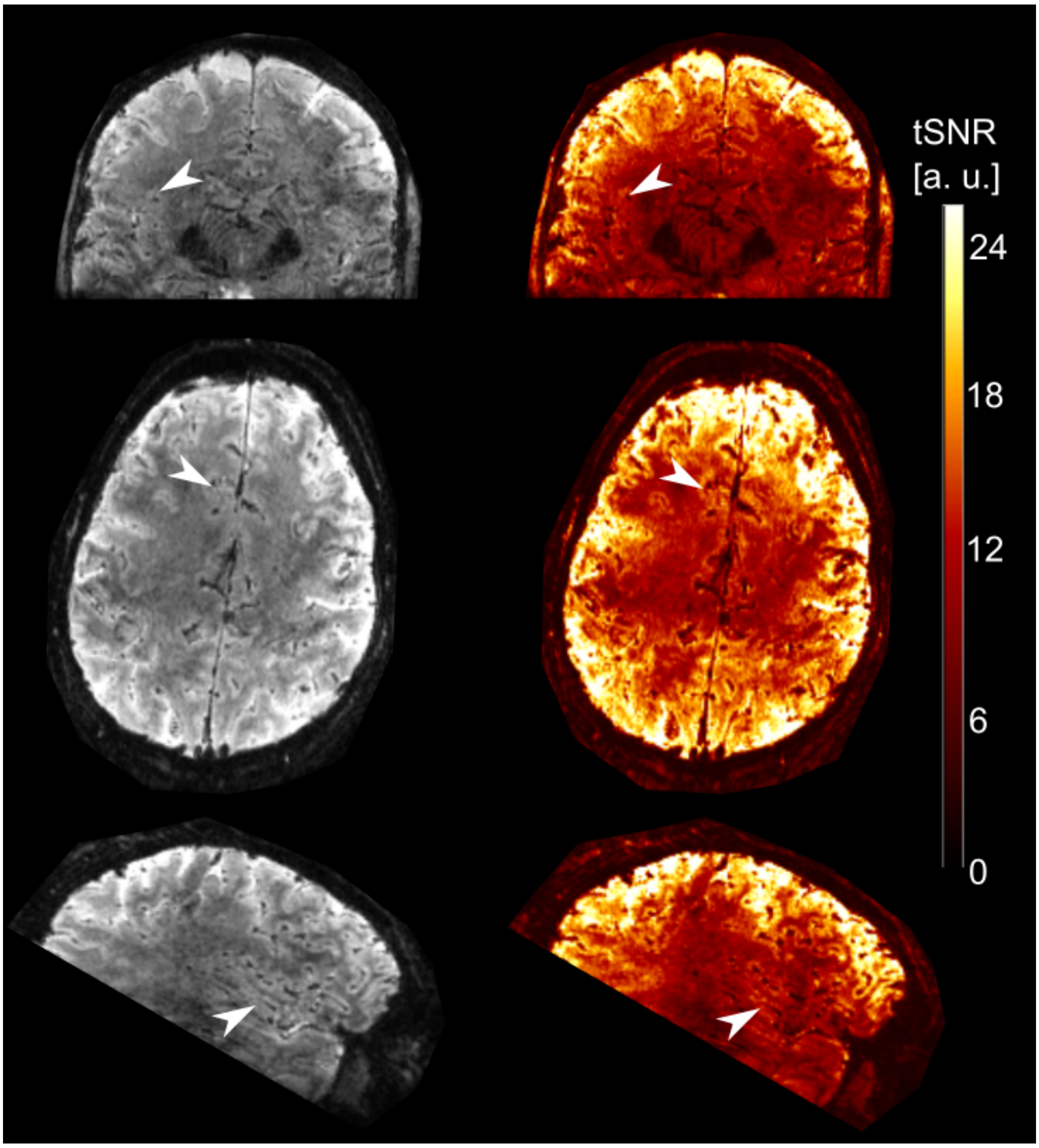


*Supplementary Figure 10 Example raw BOLD image and temporal signal-to-noise ratio.*
*Representative raw BOLD image from a pilot participant is shown together with the corresponding temporal signal-to-noise ratio (tSNR) map. The imaging slab was positioned to cover the regions of interest, including the primary and secondary somatosensory cortices and the posterior insula.*

### Supplementary tables

#### Image acquisition parameters

| Parameter | MP2RAGE | 3D-EPI |
|---|---|---|
| TR [ms] | 6000 | 54.4 |
| TE [ms] | 1.81 | 24 |
| $TR_{vol}$ [ms] | - | 3000 |
| TI1/TI2 [ms] | 800/2750 | - |
| FA1/FA2 [deg] | 4/5 | 13 |
| GRAPPA | 3 (A-P) | 4 x $2_{z1}$ ((P-A) x (H-F)) |
| No. of partitions | 192 | 108 |
| Slice direction | Sagittal | Transversal |
| FOV [mm] | 240 x 240 x 144 | 178 x 178 x 91.8 |
| Matrix size | 320 x 320 x 192 | 210 x 210 x 108 |
| Resolution [mm] | 0.75 x 0.75 x 0.75 | 0.85 x 0.85 x 0.85 |
| Phase Partial Fourier | 6/8 | 7/8 |
| Bandwidth [Hz/Px] | 290 | 1134 |
| Echo spacing [ms] | - | 1.01 |
| Acquisition time [m:s] | 14:22 | 20:50 |
| No. of volumes | 1 | 410 |
| No. of runs | 1 | 3 |

*Supplementary table 1 MRI acquisition parameters*

#### Linear mixed models with layer as a continuous variable

| Parameter | Coefficient | SE | CI | CI_low | CI_high | t | df_error | p |
|---|---|---|---|---|---|---|---|---|
| effect of BU process | 20.69 | 1.07 | 0.95 | 18.6 | 22.78 | 19.41 | 1192 | **<.001** |
| effect change by the TD process | 1.85 | 0.57 | 0.95 | 0.72 | 2.97 | 3.21 | 1192 | **<.001** |
| Linear coefficient in BU process | -1.54 | 0.27 | 0.95 | -2.08 | -1.01 | -5.71 | 1192 | **<.001** |
| Quadratic coefficient in BU process | -4.4 | 0.3 | 0.95 | -4.99 | -3.8 | -14.47 | 1192 | **<.001** |
| Linear coefficient's change in TD process (linear interaction) | 2.74 | 0.38 | 0.95 | 1.99 | 3.49 | 7.17 | 1192 | **<.001** |
| Quadratic coefficient's change in TD process (quadratic interaction) | 1.15 | 0.43 | 0.95 | 0.31 | 2 | 2.69 | 1192 | **0.01** |

*Supplementary table 2 Linear mixed-effects model parameter estimates for left S1.*
*Parameter estimates from the model fitted to data from left primary somatosensory cortex (S1) are shown. A significant main effect of process (top-down) indicates greater activation during the TD condition compared with the BU condition. The linear and quadratic terms should be interpreted jointly (see Supplementary Methods). Briefly, a negative quadratic coefficient reflects an inverted U-shaped laminar profile with a peak in middle cortical layers, whereas the linear term shifts the peak location toward deeper layers (negative coefficient) or toward superficial layers (positive coefficient). The interaction with the linear term indicates that the linear coefficient for the TD process changes sign (−1.54 + 2.74 = 1.2), and the interaction with the quadratic term indicates a reduction in curvature magnitude (−4.4 + 1.15 = −3.35). Significant interaction terms (P < 0.05) indicate that the laminar profile during TD processing differs from that during BU processing. Inspection of the parameter estimates suggests that this difference primarily involves agranular layers. This interpretation is supported by additional layer-wise interaction analyses (see Methods—Statistical analysis), which identified the marked layers shown in Fig. 2A as significant (Supplementary Tables 8–13).*

| Parameter | Coefficient | SE | CI | CI_low | CI_high | t | df_error | p |
|---|---|---|---|---|---|---|---|---|
| effect of BU process | 30.91 | 1.69 | 0.95 | 27.6 | 34.22 | 18.34 | 1192 | **<.001** |
| effect change by the TD process | 1 | 0.78 | 0.95 | -0.52 | 2.52 | 1.29 | 1192 | 0.2 |
| Linear coefficient in BU process | -1.57 | 0.37 | 0.95 | -2.28 | -0.85 | -4.29 | 1192 | **<.001** |
| Quadratic coefficient in BU process | -6.26 | 0.41 | 0.95 | -7.07 | -5.46 | -15.27 | 1192 | **<.001** |
| Linear coefficient's change in TD process (linear interaction) | 2.12 | 0.52 | 0.95 | 1.11 | 3.13 | 4.1 | 1192 | **<.001** |
| Quadratic coefficient's change in TD process (quadratic interaction) | 0.77 | 0.58 | 0.95 | -0.37 | 1.91 | 1.33 | 1192 | 0.18 |

*Supplementary table 3 Linear mixed-effects model parameter estimates for right S1.*

| Parameter | Coefficient | SE | CI | CI_low | CI_high | t | df_error | p |
|---|---|---|---|---|---|---|---|---|
| effect of BU process | 28.38 | 1.5 | 0.95 | 25.43 | 31.32 | 18.9 | 1192 | **<.001** |
| effect change by the TD process | 5.04 | 0.55 | 0.95 | 3.96 | 6.11 | 9.23 | 1192 | **<.001** |
| Linear coefficient in BU process | -1.86 | 0.26 | 0.95 | -2.37 | -1.36 | -7.26 | 1192 | **<.001** |
| Quadratic coefficient in BU process | -6.24 | 0.29 | 0.95 | -6.8 | -5.67 | -21.63 | 1192 | **<.001** |
| Linear coefficient's change in TD process (linear interaction) | 3.02 | 0.36 | 0.95 | 2.31 | 3.73 | 8.32 | 1192 | **<.001** |
| Quadratic coefficient's change in TD process (quadratic interaction) | 0.84 | 0.41 | 0.95 | 0.04 | 1.64 | 2.06 | 1192 | **0.04** |

*Supplementary table 4 Linear mixed-effects model parameter estimates for left S2.*

| Parameter | Coefficient | SE | CI | CI_low | CI_high | t | df_error | p |
|---|---|---|---|---|---|---|---|---|
| effect of BU process | 38.36 | 2.21 | 0.95 | 34.02 | 42.69 | 17.35 | 1192 | **<.001** |
| effect change by the TD process | 4.63 | 0.65 | 0.95 | 3.36 | 5.91 | 7.12 | 1192 | **<.001** |
| Linear coefficient in BU process | -0.08 | 0.31 | 0.95 | -0.68 | 0.52 | -0.25 | 1192 | 0.8 |
| Quadratic coefficient in BU process | -7.93 | 0.34 | 0.95 | -8.61 | -7.26 | -23.07 | 1192 | **<.001** |
| Linear coefficient's change in TD process (linear interaction) | 3.49 | 0.43 | 0.95 | 2.64 | 4.34 | 8.06 | 1192 | **<.001** |
| Quadratic coefficient's change in TD process (quadratic interaction) | 1.43 | 0.49 | 0.95 | 0.47 | 2.38 | 2.93 | 1192 | **<.001** |

*Supplementary table 5 Linear mixed-effects model parameter estimates for right S2.*

| Parameter | Coefficient | SE | CI | CI_low | CI_high | t | df_error | p |
|---|---|---|---|---|---|---|---|---|
| effect of BU process | 26.68 | 0.91 | 0.95 | 24.89 | 28.47 | 29.27 | 1192 | **<.001** |
| effect change by the TD process | 2.7 | 0.45 | 0.95 | 1.83 | 3.58 | 6.04 | 1192 | **<.001** |
| Linear coefficient in BU process | -2.51 | 0.21 | 0.95 | -2.92 | -2.1 | -11.9 | 1192 | **<.001** |
| Quadratic coefficient in BU process | -5.66 | 0.24 | 0.95 | -6.13 | -5.2 | -23.93 | 1192 | **<.001** |
| Linear coefficient's change in TD process (linear interaction) | 1.81 | 0.3 | 0.95 | 1.23 | 2.4 | 6.08 | 1192 | **<.001** |
| Quadratic coefficient's change in TD process (quadratic interaction) | 0.15 | 0.33 | 0.95 | -0.51 | 0.81 | 0.44 | 1192 | 0.66 |

*Supplementary table 6 Linear mixed-effects model parameter estimates for left posterior insula.*

| Parameter | Coefficient | SE | CI | CI_low | CI_high | t | df_error | p |
|---|---|---|---|---|---|---|---|---|
| effect of BU process | 29.48 | 1.27 | 0.95 | 26.99 | 31.97 | 23.24 | 1192 | **<.001** |
| effect change by the TD process | 0.47 | 0.51 | 0.95 | -0.53 | 1.47 | 0.93 | 1192 | 0.35 |
| Linear coefficient in BU process | -0.47 | 0.24 | 0.95 | -0.93 | 0 | -1.95 | 1192 | **0.05** |
| Quadratic coefficient in BU process | -6.1 | 0.27 | 0.95 | -6.63 | -5.58 | -22.74 | 1192 | **<.001** |
| Linear coefficient's change in TD process (linear interaction) | 0.8 | 0.34 | 0.95 | 0.13 | 1.46 | 2.35 | 1192 | **0.02** |
| Quadratic coefficient's change in TD process (quadratic interaction) | 0.53 | 0.38 | 0.95 | -0.22 | 1.27 | 1.38 | 1192 | 0.17 |

*Supplementary table 7 Linear mixed-effects model parameter estimates for right posterior insula.*

### Linear mixed models with layer as a factor variable

| Parameter | Coefficient | SE | CI | CI_low | CI_high | t | df_error | p | p_fdr |
|---|---|---|---|---|---|---|---|---|---|
| effect of BU process | 11.28 | 1.57 | 0.95 | 8.19 | 14.36 | 7.17 | 1158 | <.001 | NA |
| effect change by the TD process | 0.45 | 1.73 | 0.95 | -2.95 | 3.85 | 0.26 | 1158 | 0.8 | NA |
| effect change by layer 2 in BU | 1.94 | 1.73 | 0.95 | -1.46 | 5.34 | 1.12 | 1158 | 0.26 | NA |
| effect change by layer 3 in BU | 3.81 | 1.73 | 0.95 | 0.41 | 7.22 | 2.2 | 1158 | 0.03 | NA |
| effect change by layer 4 in BU | 5.54 | 1.73 | 0.95 | 2.14 | 8.94 | 3.2 | 1158 | <.001 | NA |
| effect change by layer 5 in BU | 6.99 | 1.73 | 0.95 | 3.58 | 10.39 | 4.03 | 1158 | <.001 | NA |
| effect change by layer 6 in BU | 8.11 | 1.73 | 0.95 | 4.7 | 11.51 | 4.68 | 1158 | <.001 | NA |
| effect change by layer 7 in BU | 8.91 | 1.73 | 0.95 | 5.51 | 12.31 | 5.14 | 1158 | <.001 | NA |
| effect change by layer 8 in BU | 9.43 | 1.73 | 0.95 | 6.02 | 12.83 | 5.44 | 1158 | <.001 | NA |
| effect change by layer 9 in BU | 9.66 | 1.73 | 0.95 | 6.26 | 13.06 | 5.57 | 1158 | <.001 | NA |
| effect change by layer 10 in BU | 9.63 | 1.73 | 0.95 | 6.22 | 13.03 | 5.55 | 1158 | <.001 | NA |
| effect change by layer 11 in BU | 9.36 | 1.73 | 0.95 | 5.96 | 12.76 | 5.4 | 1158 | <.001 | NA |
| effect change by layer 12 in BU | 8.85 | 1.73 | 0.95 | 5.45 | 12.25 | 5.11 | 1158 | <.001 | NA |
| effect change by layer 13 in BU | 8.04 | 1.73 | 0.95 | 4.64 | 11.44 | 4.64 | 1158 | <.001 | NA |
| effect change by layer 14 in BU | 6.9 | 1.73 | 0.95 | 3.5 | 10.3 | 3.98 | 1158 | <.001 | NA |
| effect change by layer 15 in BU | 5.37 | 1.73 | 0.95 | 1.97 | 8.77 | 3.1 | 1158 | <.001 | NA |
| effect change by layer 16 in BU | 3.47 | 1.73 | 0.95 | 0.07 | 6.87 | 2 | 1158 | 0.05 | NA |
| effect change by layer 17 in BU | 1.42 | 1.73 | 0.95 | -1.98 | 4.82 | 0.82 | 1158 | 0.41 | NA |
| effect change by layer 18 in BU | -0.59 | 1.73 | 0.95 | -3.99 | 2.81 | -0.34 | 1158 | 0.73 | NA |
| effect change by layer 19 in BU | -2.41 | 1.73 | 0.95 | -5.81 | 0.99 | -1.39 | 1158 | 0.16 | NA |
| effect change by layer 20 in BU | -3.98 | 1.73 | 0.95 | -7.38 | -0.58 | -2.3 | 1158 | 0.02 | NA |
| change of layer 2 effect by TD | 0 | 2.45 | 0.95 | -4.81 | 4.81 | 0 | 1158 | 1 | 1 |
| change of layer 3 effect by TD | -0.05 | 2.45 | 0.95 | -4.86 | 4.76 | -0.02 | 1158 | 0.98 | 1 |
| change of layer 4 effect by TD | -0.16 | 2.45 | 0.95 | -4.97 | 4.65 | -0.07 | 1158 | 0.95 | 1 |
| change of layer 5 effect by TD | -0.24 | 2.45 | 0.95 | -5.05 | 4.57 | -0.1 | 1158 | 0.92 | 1 |
| change of layer 6 effect by TD | -0.24 | 2.45 | 0.95 | -5.05 | 4.57 | -0.1 | 1158 | 0.92 | 1 |
| change of layer 7 effect by TD | -0.12 | 2.45 | 0.95 | -4.93 | 4.69 | -0.05 | 1158 | 0.96 | 1 |
| change of layer 8 effect by TD | 0.2 | 2.45 | 0.95 | -4.61 | 5.01 | 0.08 | 1158 | 0.94 | 1 |
| change of layer 9 effect by TD | 0.67 | 2.45 | 0.95 | -4.14 | 5.48 | 0.28 | 1158 | 0.78 | 1 |
| change of layer 10 effect by TD | 1.24 | 2.45 | 0.95 | -3.56 | 6.05 | 0.51 | 1158 | 0.61 | 1 |
| change of layer 11 effect by TD | 1.83 | 2.45 | 0.95 | -2.98 | 6.64 | 0.74 | 1158 | 0.46 | 0.87 |
| change of layer 12 effect by TD | 2.4 | 2.45 | 0.95 | -2.41 | 7.21 | 0.98 | 1158 | 0.33 | 0.69 |
| change of layer 13 effect by TD | 2.91 | 2.45 | 0.95 | -1.9 | 7.72 | 1.19 | 1158 | 0.23 | 0.56 |
| change of layer 14 effect by TD | 3.45 | 2.45 | 0.95 | -1.36 | 8.26 | 1.41 | 1158 | 0.16 | 0.43 |
| change of layer 15 effect by TD | 4.16 | 2.45 | 0.95 | -0.65 | 8.97 | 1.7 | 1158 | 0.09 | 0.28 |
| change of layer 16 effect by TD | 5.08 | 2.45 | 0.95 | 0.27 | 9.89 | 2.07 | 1158 | **0.04** | 0.15 |
| change of layer 17 effect by TD | 6.06 | 2.45 | 0.95 | 1.25 | 10.87 | 2.47 | 1158 | **0.01** | 0.06 |
| change of layer 18 effect by TD | 7.06 | 2.45 | 0.95 | 2.25 | 11.87 | 2.88 | 1158 | **<.001** | **0.03** |
| change of layer 19 effect by TD | 7.98 | 2.45 | 0.95 | 3.17 | 12.79 | 3.25 | 1158 | **<.001** | **0.01** |
| change of layer 20 effect by TD | 8.77 | 2.45 | 0.95 | 3.96 | 13.58 | 3.58 | 1158 | **<.001** | **0.01** |

*Supplementary table 8 Robust linear mixed-effects model parameter estimates for left S1.*
*Parameter estimates from a robust linear mixed-effects model fitted to data from left primary somatosensory cortex (S1) are reported. P values < 0.05 are shown in bold. Layer was modeled as a categorical (factor) variable, such that each layer was tested relative to the reference level, defined as the bottom-up (BU) process in the deepest layer (layer 1). Layer indices increase from deep to superficial (layers 1–20).*
*“Effect change by layer in BU” denotes the coefficient estimate for each layer relative to the reference level, whereas “change of layer effect by TD” denotes the interaction term, reflecting how the layer-specific effect is modulated by the top-down (TD) process. False discovery rate (FDR) correction was applied to p values to control for type I error. P-values below significant threshold are highlighted in bold in the interesting contrasts (interaction effect).*

| Parameter | Coefficient | SE | CI | CI_low | CI_high | t | df_error | p | p_fdr |
|---|---|---|---|---|---|---|---|---|---|
| effect of BU process | 16.83 | 2.3 | 0.95 | 12.33 | 21.34 | 7.33 | 1158 | <.001 | NA |
| effect change by the TD process | -0.18 | 2.34 | 0.95 | -4.77 | 4.4 | -0.08 | 1158 | 0.94 | NA |
| effect change by layer 2 in BU | 2.74 | 2.34 | 0.95 | -1.84 | 7.33 | 1.17 | 1158 | 0.24 | NA |
| effect change by layer 3 in BU | 5.34 | 2.34 | 0.95 | 0.75 | 9.93 | 2.28 | 1158 | 0.02 | NA |
| effect change by layer 4 in BU | 7.63 | 2.34 | 0.95 | 3.04 | 12.21 | 3.26 | 1158 | <.001 | NA |
| effect change by layer 5 in BU | 9.58 | 2.34 | 0.95 | 4.99 | 14.16 | 4.1 | 1158 | <.001 | NA |
| effect change by layer 6 in BU | 11.23 | 2.34 | 0.95 | 6.64 | 15.82 | 4.8 | 1158 | <.001 | NA |
| effect change by layer 7 in BU | 12.58 | 2.34 | 0.95 | 8 | 17.17 | 5.38 | 1158 | <.001 | NA |
| effect change by layer 8 in BU | 13.61 | 2.34 | 0.95 | 9.03 | 18.2 | 5.82 | 1158 | <.001 | NA |
| effect change by layer 9 in BU | 14.34 | 2.34 | 0.95 | 9.75 | 18.92 | 6.13 | 1158 | <.001 | NA |
| effect change by layer 10 in BU | 14.64 | 2.34 | 0.95 | 10.06 | 19.23 | 6.26 | 1158 | <.001 | NA |
| effect change by layer 11 in BU | 14.51 | 2.34 | 0.95 | 9.92 | 19.1 | 6.21 | 1158 | <.001 | NA |
| effect change by layer 12 in BU | 13.87 | 2.34 | 0.95 | 9.28 | 18.46 | 5.93 | 1158 | <.001 | NA |
| effect change by layer 13 in BU | 12.66 | 2.34 | 0.95 | 8.08 | 17.25 | 5.42 | 1158 | <.001 | NA |
| effect change by layer 14 in BU | 10.93 | 2.34 | 0.95 | 6.34 | 15.52 | 4.67 | 1158 | <.001 | NA |
| effect change by layer 15 in BU | 8.67 | 2.34 | 0.95 | 4.09 | 13.26 | 3.71 | 1158 | <.001 | NA |
| effect change by layer 16 in BU | 6.05 | 2.34 | 0.95 | 1.46 | 10.64 | 2.59 | 1158 | 0.01 | NA |
| effect change by layer 17 in BU | 3.29 | 2.34 | 0.95 | -1.3 | 7.87 | 1.41 | 1158 | 0.16 | NA |
| effect change by layer 18 in BU | 0.57 | 2.34 | 0.95 | -4.02 | 5.16 | 0.24 | 1158 | 0.81 | NA |
| effect change by layer 19 in BU | -1.9 | 2.34 | 0.95 | -6.49 | 2.69 | -0.81 | 1158 | 0.42 | NA |
| effect change by layer 20 in BU | -3.95 | 2.34 | 0.95 | -8.54 | 0.64 | -1.69 | 1158 | 0.09 | NA |
| change of layer 2 effect by TD | -0.05 | 3.31 | 0.95 | -6.54 | 6.44 | -0.02 | 1158 | 0.99 | 0.99 |
| change of layer 3 effect by TD | -0.17 | 3.31 | 0.95 | -6.66 | 6.31 | -0.05 | 1158 | 0.96 | 0.99 |
| change of layer 4 effect by TD | -0.27 | 3.31 | 0.95 | -6.76 | 6.22 | -0.08 | 1158 | 0.93 | 0.99 |
| change of layer 5 effect by TD | -0.31 | 3.31 | 0.95 | -6.8 | 6.17 | -0.09 | 1158 | 0.92 | 0.99 |
| change of layer 6 effect by TD | -0.3 | 3.31 | 0.95 | -6.78 | 6.19 | -0.09 | 1158 | 0.93 | 0.99 |
| change of layer 7 effect by TD | -0.16 | 3.31 | 0.95 | -6.65 | 6.32 | -0.05 | 1158 | 0.96 | 0.99 |
| change of layer 8 effect by TD | 0.09 | 3.31 | 0.95 | -6.39 | 6.58 | 0.03 | 1158 | 0.98 | 0.99 |
| change of layer 9 effect by TD | 0.47 | 3.31 | 0.95 | -6.02 | 6.95 | 0.14 | 1158 | 0.89 | 0.99 |
| change of layer 10 effect by TD | 0.99 | 3.31 | 0.95 | -5.5 | 7.47 | 0.3 | 1158 | 0.77 | 0.99 |
| change of layer 11 effect by TD | 1.59 | 3.31 | 0.95 | -4.9 | 8.07 | 0.48 | 1158 | 0.63 | 0.99 |
| change of layer 12 effect by TD | 2.14 | 3.31 | 0.95 | -4.35 | 8.62 | 0.65 | 1158 | 0.52 | 0.99 |
| change of layer 13 effect by TD | 2.65 | 3.31 | 0.95 | -3.84 | 9.14 | 0.8 | 1158 | 0.42 | 0.99 |
| change of layer 14 effect by TD | 3.04 | 3.31 | 0.95 | -3.45 | 9.52 | 0.92 | 1158 | 0.36 | 0.97 |
| change of layer 15 effect by TD | 3.44 | 3.31 | 0.95 | -3.05 | 9.93 | 1.04 | 1158 | 0.3 | 0.95 |
| change of layer 16 effect by TD | 3.89 | 3.31 | 0.95 | -2.59 | 10.38 | 1.18 | 1158 | 0.24 | 0.91 |
| change of layer 17 effect by TD | 4.46 | 3.31 | 0.95 | -2.03 | 10.95 | 1.35 | 1158 | 0.18 | 0.84 |
| change of layer 18 effect by TD | 5.11 | 3.31 | 0.95 | -1.38 | 11.6 | 1.55 | 1158 | 0.12 | 0.78 |
| change of layer 19 effect by TD | 5.9 | 3.31 | 0.95 | -0.59 | 12.38 | 1.78 | 1158 | 0.07 | 0.71 |
| change of layer 20 effect by TD | 6.65 | 3.31 | 0.95 | 0.16 | 13.14 | 2.01 | 1158 | **0.04** | 0.71 |

*Supplementary table 9 Robust linear mixed-effects model parameter estimates for right S1.*

| Parameter | Coefficient | SE | CI | CI_low | CI_high | t | df_error | p | p_fdr |
|---|---|---|---|---|---|---|---|---|---|
| effect of BU process | 14.19 | 1.86 | 0.95 | 10.54 | 17.83 | 7.64 | 1158 | <.001 | NA |
| effect change by the TD process | 2.86 | 1.64 | 0.95 | -0.36 | 6.07 | 1.74 | 1158 | 0.08 | NA |
| effect change by layer 2 in BU | 3.1 | 1.64 | 0.95 | -0.12 | 6.31 | 1.89 | 1158 | 0.06 | NA |
| effect change by layer 3 in BU | 5.99 | 1.64 | 0.95 | 2.77 | 9.2 | 3.66 | 1158 | <.001 | NA |
| effect change by layer 4 in BU | 8.46 | 1.64 | 0.95 | 5.25 | 11.68 | 5.17 | 1158 | <.001 | NA |
| effect change by layer 5 in BU | 10.45 | 1.64 | 0.95 | 7.23 | 13.66 | 6.38 | 1158 | <.001 | NA |
| effect change by layer 6 in BU | 12 | 1.64 | 0.95 | 8.79 | 15.21 | 7.33 | 1158 | <.001 | NA |
| effect change by layer 7 in BU | 13.15 | 1.64 | 0.95 | 9.94 | 16.36 | 8.03 | 1158 | <.001 | NA |
| effect change by layer 8 in BU | 14 | 1.64 | 0.95 | 10.79 | 17.21 | 8.55 | 1158 | <.001 | NA |
| effect change by layer 9 in BU | 14.55 | 1.64 | 0.95 | 11.34 | 17.77 | 8.89 | 1158 | <.001 | NA |
| effect change by layer 10 in BU | 14.7 | 1.64 | 0.95 | 11.49 | 17.91 | 8.98 | 1158 | <.001 | NA |
| effect change by layer 11 in BU | 14.49 | 1.64 | 0.95 | 11.27 | 17.7 | 8.85 | 1158 | <.001 | NA |
| effect change by layer 12 in BU | 13.82 | 1.64 | 0.95 | 10.6 | 17.03 | 8.44 | 1158 | <.001 | NA |
| effect change by layer 13 in BU | 12.62 | 1.64 | 0.95 | 9.41 | 15.84 | 7.71 | 1158 | <.001 | NA |
| effect change by layer 14 in BU | 10.76 | 1.64 | 0.95 | 7.55 | 13.98 | 6.57 | 1158 | <.001 | NA |
| effect change by layer 15 in BU | 8.31 | 1.64 | 0.95 | 5.09 | 11.52 | 5.07 | 1158 | <.001 | NA |
| effect change by layer 16 in BU | 5.47 | 1.64 | 0.95 | 2.25 | 8.68 | 3.34 | 1158 | <.001 | NA |
| effect change by layer 17 in BU | 2.63 | 1.64 | 0.95 | -0.58 | 5.84 | 1.61 | 1158 | 0.11 | NA |
| effect change by layer 18 in BU | 0.14 | 1.64 | 0.95 | -3.08 | 3.35 | 0.08 | 1158 | 0.93 | NA |
| effect change by layer 19 in BU | -1.93 | 1.64 | 0.95 | -5.14 | 1.29 | -1.18 | 1158 | 0.24 | NA |
| effect change by layer 20 in BU | -3.52 | 1.64 | 0.95 | -6.73 | -0.31 | -2.15 | 1158 | 0.03 | NA |
| change of layer 2 effect by TD | 0.06 | 2.32 | 0.95 | -4.48 | 4.6 | 0.03 | 1158 | 0.98 | 0.99 |
| change of layer 3 effect by TD | -0.03 | 2.32 | 0.95 | -4.57 | 4.52 | -0.01 | 1158 | 0.99 | 0.99 |
| change of layer 4 effect by TD | -0.21 | 2.32 | 0.95 | -4.75 | 4.33 | -0.09 | 1158 | 0.93 | 0.99 |
| change of layer 5 effect by TD | -0.32 | 2.32 | 0.95 | -4.87 | 4.22 | -0.14 | 1158 | 0.89 | 0.99 |
| change of layer 6 effect by TD | -0.31 | 2.32 | 0.95 | -4.86 | 4.23 | -0.13 | 1158 | 0.89 | 0.99 |
| change of layer 7 effect by TD | -0.1 | 2.32 | 0.95 | -4.65 | 4.44 | -0.04 | 1158 | 0.96 | 0.99 |
| change of layer 8 effect by TD | 0.32 | 2.32 | 0.95 | -4.22 | 4.86 | 0.14 | 1158 | 0.89 | 0.99 |
| change of layer 9 effect by TD | 0.93 | 2.32 | 0.95 | -3.61 | 5.47 | 0.4 | 1158 | 0.69 | 0.99 |
| change of layer 10 effect by TD | 1.74 | 2.32 | 0.95 | -2.81 | 6.28 | 0.75 | 1158 | 0.45 | 0.78 |
| change of layer 11 effect by TD | 2.55 | 2.32 | 0.95 | -1.99 | 7.09 | 1.1 | 1158 | 0.27 | 0.52 |
| change of layer 12 effect by TD | 3.33 | 2.32 | 0.95 | -1.22 | 7.87 | 1.44 | 1158 | 0.15 | 0.32 |
| change of layer 13 effect by TD | 4.1 | 2.32 | 0.95 | -0.44 | 8.65 | 1.77 | 1158 | 0.08 | 0.18 |
| change of layer 14 effect by TD | 4.89 | 2.32 | 0.95 | 0.35 | 9.44 | 2.11 | 1158 | **0.03** | 0.09 |
| change of layer 15 effect by TD | 5.71 | 2.32 | 0.95 | 1.17 | 10.25 | 2.47 | 1158 | **0.01** | **0.04** |
| change of layer 16 effect by TD | 6.54 | 2.32 | 0.95 | 2 | 11.08 | 2.82 | 1158 | **<.001** | **0.02** |
| change of layer 17 effect by TD | 7.2 | 2.32 | 0.95 | 2.65 | 11.74 | 3.11 | 1158 | **<.001** | **0.01** |
| change of layer 18 effect by TD | 7.69 | 2.32 | 0.95 | 3.15 | 12.24 | 3.32 | 1158 | **<.001** | **0.01** |
| change of layer 19 effect by TD | 8.05 | 2.32 | 0.95 | 3.5 | 12.59 | 3.47 | 1158 | **<.001** | **0.01** |
| change of layer 20 effect by TD | 8.26 | 2.32 | 0.95 | 3.72 | 12.81 | 3.57 | 1158 | **<.001** | **0.01** |

*Supplementary table 10 Robust linear mixed-effects model parameter estimates for left S2.*

| Parameter | Coefficient | SE | CI | CI_low | CI_high | t | df_error | p | p_fdr |
|---|---|---|---|---|---|---|---|---|---|
| effect of BU process | 18.47 | 2.56 | 0.95 | 13.45 | 23.5 | 7.21 | 1158 | <.001 | NA |
| effect change by the TD process | 2.27 | 1.94 | 0.95 | -1.54 | 6.08 | 1.17 | 1158 | 0.24 | NA |
| effect change by layer 2 in BU | 3.5 | 1.94 | 0.95 | -0.31 | 7.31 | 1.8 | 1158 | 0.07 | NA |
| effect change by layer 3 in BU | 6.65 | 1.94 | 0.95 | 2.84 | 10.46 | 3.42 | 1158 | <.001 | NA |
| effect change by layer 4 in BU | 9.32 | 1.94 | 0.95 | 5.51 | 13.13 | 4.8 | 1158 | <.001 | NA |
| effect change by layer 5 in BU | 11.61 | 1.94 | 0.95 | 7.8 | 15.42 | 5.98 | 1158 | <.001 | NA |
| effect change by layer 6 in BU | 13.62 | 1.94 | 0.95 | 9.81 | 17.43 | 7.02 | 1158 | <.001 | NA |
| effect change by layer 7 in BU | 15.46 | 1.94 | 0.95 | 11.65 | 19.27 | 7.96 | 1158 | <.001 | NA |
| effect change by layer 8 in BU | 17.19 | 1.94 | 0.95 | 13.38 | 21 | 8.85 | 1158 | <.001 | NA |
| effect change by layer 9 in BU | 18.83 | 1.94 | 0.95 | 15.02 | 22.64 | 9.7 | 1158 | <.001 | NA |
| effect change by layer 10 in BU | 20.26 | 1.94 | 0.95 | 16.45 | 24.07 | 10.43 | 1158 | <.001 | NA |
| effect change by layer 11 in BU | 21.2 | 1.94 | 0.95 | 17.39 | 25.01 | 10.92 | 1158 | <.001 | NA |
| effect change by layer 12 in BU | 21.3 | 1.94 | 0.95 | 17.49 | 25.11 | 10.97 | 1158 | <.001 | NA |
| effect change by layer 13 in BU | 20.36 | 1.94 | 0.95 | 16.55 | 24.17 | 10.48 | 1158 | <.001 | NA |
| effect change by layer 14 in BU | 18.36 | 1.94 | 0.95 | 14.55 | 22.17 | 9.45 | 1158 | <.001 | NA |
| effect change by layer 15 in BU | 15.43 | 1.94 | 0.95 | 11.62 | 19.24 | 7.95 | 1158 | <.001 | NA |
| effect change by layer 16 in BU | 11.89 | 1.94 | 0.95 | 8.08 | 15.7 | 6.12 | 1158 | <.001 | NA |
| effect change by layer 17 in BU | 8.15 | 1.94 | 0.95 | 4.34 | 11.96 | 4.2 | 1158 | <.001 | NA |
| effect change by layer 18 in BU | 4.72 | 1.94 | 0.95 | 0.91 | 8.53 | 2.43 | 1158 | 0.02 | NA |
| effect change by layer 19 in BU | 1.82 | 1.94 | 0.95 | -1.99 | 5.63 | 0.94 | 1158 | 0.35 | NA |
| effect change by layer 20 in BU | -0.47 | 1.94 | 0.95 | -4.28 | 3.34 | -0.24 | 1158 | 0.81 | NA |
| change of layer 2 effect by TD | 0.31 | 2.75 | 0.95 | -5.08 | 5.7 | 0.11 | 1158 | 0.91 | 0.91 |
| change of layer 3 effect by TD | 0.58 | 2.75 | 0.95 | -4.8 | 5.97 | 0.21 | 1158 | 0.83 | 0.88 |
| change of layer 4 effect by TD | 0.74 | 2.75 | 0.95 | -4.65 | 6.13 | 0.27 | 1158 | 0.79 | 0.88 |
| change of layer 5 effect by TD | 0.74 | 2.75 | 0.95 | -4.65 | 6.13 | 0.27 | 1158 | 0.79 | 0.88 |
| change of layer 6 effect by TD | 0.74 | 2.75 | 0.95 | -4.65 | 6.13 | 0.27 | 1158 | 0.79 | 0.88 |
| change of layer 7 effect by TD | 0.76 | 2.75 | 0.95 | -4.63 | 6.15 | 0.28 | 1158 | 0.78 | 0.88 |
| change of layer 8 effect by TD | 0.93 | 2.75 | 0.95 | -4.46 | 6.32 | 0.34 | 1158 | 0.73 | 0.88 |
| change of layer 9 effect by TD | 1.22 | 2.75 | 0.95 | -4.16 | 6.61 | 0.45 | 1158 | 0.66 | 0.88 |
| change of layer 10 effect by TD | 1.5 | 2.75 | 0.95 | -3.89 | 6.89 | 0.55 | 1158 | 0.58 | 0.88 |
| change of layer 11 effect by TD | 1.98 | 2.75 | 0.95 | -3.41 | 7.37 | 0.72 | 1158 | 0.47 | 0.88 |
| change of layer 12 effect by TD | 2.77 | 2.75 | 0.95 | -2.62 | 8.16 | 1.01 | 1158 | 0.31 | 0.66 |
| change of layer 13 effect by TD | 3.84 | 2.75 | 0.95 | -1.54 | 9.23 | 1.4 | 1158 | 0.16 | 0.38 |
| change of layer 14 effect by TD | 5.2 | 2.75 | 0.95 | -0.19 | 10.58 | 1.89 | 1158 | 0.06 | 0.16 |
| change of layer 15 effect by TD | 6.59 | 2.75 | 0.95 | 1.2 | 11.98 | 2.4 | 1158 | **0.02** | **0.05** |
| change of layer 16 effect by TD | 7.84 | 2.75 | 0.95 | 2.46 | 13.23 | 2.86 | 1158 | **<.001** | **0.02** |
| change of layer 17 effect by TD | 8.96 | 2.75 | 0.95 | 3.57 | 14.35 | 3.26 | 1158 | **<.001** | **0.01** |
| change of layer 18 effect by TD | 9.79 | 2.75 | 0.95 | 4.4 | 15.17 | 3.56 | 1158 | **<.001** | **<.001** |
| change of layer 19 effect by TD | 10.4 | 2.75 | 0.95 | 5.01 | 15.79 | 3.79 | 1158 | **<.001** | **<.001** |
| change of layer 20 effect by TD | 10.92 | 2.75 | 0.95 | 5.53 | 16.3 | 3.97 | 1158 | **<.001** | **<.001** |

*Supplementary table 11 Robust linear mixed-effects model parameter estimates for right S2.*

| Parameter | Coefficient | SE | CI | CI_low | CI_high | t | df_error | p | p_fdr |
|---|---|---|---|---|---|---|---|---|---|
| effect of BU process | 14.79 | 1.27 | 0.95 | 12.29 | 17.29 | 11.6 | 1158 | <.001 | NA |
| effect change by the TD process | 0.8 | 1.34 | 0.95 | -1.82 | 3.43 | 0.6 | 1158 | 0.55 | NA |
| effect change by layer 2 in BU | 3 | 1.34 | 0.95 | 0.37 | 5.62 | 2.24 | 1158 | 0.03 | NA |
| effect change by layer 3 in BU | 5.7 | 1.34 | 0.95 | 3.07 | 8.32 | 4.26 | 1158 | <.001 | NA |
| effect change by layer 4 in BU | 7.89 | 1.34 | 0.95 | 5.26 | 10.51 | 5.9 | 1158 | <.001 | NA |
| effect change by layer 5 in BU | 9.47 | 1.34 | 0.95 | 6.85 | 12.09 | 7.08 | 1158 | <.001 | NA |
| effect change by layer 6 in BU | 10.6 | 1.34 | 0.95 | 7.98 | 13.23 | 7.93 | 1158 | <.001 | NA |
| effect change by layer 7 in BU | 11.39 | 1.34 | 0.95 | 8.76 | 14.01 | 8.52 | 1158 | <.001 | NA |
| effect change by layer 8 in BU | 11.98 | 1.34 | 0.95 | 9.36 | 14.6 | 8.96 | 1158 | <.001 | NA |
| effect change by layer 9 in BU | 12.35 | 1.34 | 0.95 | 9.73 | 14.97 | 9.23 | 1158 | <.001 | NA |
| effect change by layer 10 in BU | 12.38 | 1.34 | 0.95 | 9.76 | 15.01 | 9.26 | 1158 | <.001 | NA |
| effect change by layer 11 in BU | 12.08 | 1.34 | 0.95 | 9.46 | 14.71 | 9.04 | 1158 | <.001 | NA |
| effect change by layer 12 in BU | 11.33 | 1.34 | 0.95 | 8.7 | 13.95 | 8.47 | 1158 | <.001 | NA |
| effect change by layer 13 in BU | 10.02 | 1.34 | 0.95 | 7.4 | 12.65 | 7.5 | 1158 | <.001 | NA |
| effect change by layer 14 in BU | 8.18 | 1.34 | 0.95 | 5.55 | 10.8 | 6.11 | 1158 | <.001 | NA |
| effect change by layer 15 in BU | 5.86 | 1.34 | 0.95 | 3.24 | 8.48 | 4.38 | 1158 | <.001 | NA |
| effect change by layer 16 in BU | 3.26 | 1.34 | 0.95 | 0.63 | 5.88 | 2.44 | 1158 | 0.02 | NA |
| effect change by layer 17 in BU | 0.6 | 1.34 | 0.95 | -2.03 | 3.22 | 0.45 | 1158 | 0.66 | NA |
| effect change by layer 18 in BU | -1.84 | 1.34 | 0.95 | -4.46 | 0.79 | -1.37 | 1158 | 0.17 | NA |
| effect change by layer 19 in BU | -3.94 | 1.34 | 0.95 | -6.56 | -1.31 | -2.94 | 1158 | <.001 | NA |
| effect change by layer 20 in BU | -5.59 | 1.34 | 0.95 | -8.21 | -2.96 | -4.18 | 1158 | <.001 | NA |
| change of layer 2 effect by TD | -0.12 | 1.89 | 0.95 | -3.83 | 3.59 | -0.06 | 1158 | 0.95 | 0.97 |
| change of layer 3 effect by TD | -0.27 | 1.89 | 0.95 | -3.98 | 3.44 | -0.14 | 1158 | 0.89 | 0.97 |
| change of layer 4 effect by TD | -0.42 | 1.89 | 0.95 | -4.13 | 3.29 | -0.22 | 1158 | 0.83 | 0.97 |
| change of layer 5 effect by TD | -0.35 | 1.89 | 0.95 | -4.06 | 3.36 | -0.19 | 1158 | 0.85 | 0.97 |
| change of layer 6 effect by TD | -0.08 | 1.89 | 0.95 | -3.79 | 3.63 | -0.04 | 1158 | 0.97 | 0.97 |
| change of layer 7 effect by TD | 0.4 | 1.89 | 0.95 | -3.31 | 4.11 | 0.21 | 1158 | 0.83 | 0.97 |
| change of layer 8 effect by TD | 1.01 | 1.89 | 0.95 | -2.7 | 4.72 | 0.53 | 1158 | 0.59 | 0.87 |
| change of layer 9 effect by TD | 1.58 | 1.89 | 0.95 | -2.13 | 5.29 | 0.83 | 1158 | 0.4 | 0.64 |
| change of layer 10 effect by TD | 2.15 | 1.89 | 0.95 | -1.56 | 5.86 | 1.14 | 1158 | 0.26 | 0.44 |
| change of layer 11 effect by TD | 2.6 | 1.89 | 0.95 | -1.11 | 6.31 | 1.38 | 1158 | 0.17 | 0.32 |
| change of layer 12 effect by TD | 2.93 | 1.89 | 0.95 | -0.78 | 6.64 | 1.55 | 1158 | 0.12 | 0.26 |
| change of layer 13 effect by TD | 3.14 | 1.89 | 0.95 | -0.57 | 6.85 | 1.66 | 1158 | 0.1 | 0.23 |
| change of layer 14 effect by TD | 3.26 | 1.89 | 0.95 | -0.45 | 6.97 | 1.72 | 1158 | 0.09 | 0.23 |
| change of layer 15 effect by TD | 3.31 | 1.89 | 0.95 | -0.4 | 7.02 | 1.75 | 1158 | 0.08 | 0.23 |
| change of layer 16 effect by TD | 3.48 | 1.89 | 0.95 | -0.23 | 7.19 | 1.84 | 1158 | 0.07 | 0.23 |
| change of layer 17 effect by TD | 3.82 | 1.89 | 0.95 | 0.11 | 7.53 | 2.02 | 1158 | **0.04** | 0.21 |
| change of layer 18 effect by TD | 4.28 | 1.89 | 0.95 | 0.57 | 7.99 | 2.27 | 1158 | **0.02** | 0.15 |
| change of layer 19 effect by TD | 4.86 | 1.89 | 0.95 | 1.15 | 8.57 | 2.57 | 1158 | **0.01** | 0.1 |
| change of layer 20 effect by TD | 5.38 | 1.89 | 0.95 | 1.67 | 9.09 | 2.85 | 1158 | **<.001** | 0.09 |

*Supplementary table 12 Robust linear mixed-effects model parameter estimates for left posterior insula.*

| Parameter | Coefficient | SE | CI | CI_low | CI_high | t | df_error | p | p_fdr |
|---|---|---|---|---|---|---|---|---|---|
| effect of BU process | 13.91 | 1.63 | 0.95 | 10.73 | 17.1 | 8.56 | 1158 | <.001 | NA |
| effect change by the TD process | 0.68 | 1.52 | 0.95 | -2.31 | 3.67 | 0.45 | 1158 | 0.66 | NA |
| effect change by layer 2 in BU | 3.1 | 1.52 | 0.95 | 0.11 | 6.09 | 2.03 | 1158 | 0.04 | NA |
| effect change by layer 3 in BU | 5.91 | 1.52 | 0.95 | 2.91 | 8.9 | 3.87 | 1158 | <.001 | NA |
| effect change by layer 4 in BU | 8.26 | 1.52 | 0.95 | 5.27 | 11.25 | 5.42 | 1158 | <.001 | NA |
| effect change by layer 5 in BU | 10.21 | 1.52 | 0.95 | 7.22 | 13.2 | 6.7 | 1158 | <.001 | NA |
| effect change by layer 6 in BU | 11.84 | 1.52 | 0.95 | 8.85 | 14.83 | 7.77 | 1158 | <.001 | NA |
| effect change by layer 7 in BU | 13.19 | 1.52 | 0.95 | 10.19 | 16.18 | 8.65 | 1158 | <.001 | NA |
| effect change by layer 8 in BU | 14.32 | 1.52 | 0.95 | 11.33 | 17.31 | 9.39 | 1158 | <.001 | NA |
| effect change by layer 9 in BU | 15.24 | 1.52 | 0.95 | 12.25 | 18.23 | 9.99 | 1158 | <.001 | NA |
| effect change by layer 10 in BU | 15.8 | 1.52 | 0.95 | 12.81 | 18.79 | 10.36 | 1158 | <.001 | NA |
| effect change by layer 11 in BU | 16.03 | 1.52 | 0.95 | 13.04 | 19.02 | 10.52 | 1158 | <.001 | NA |
| effect change by layer 12 in BU | 15.8 | 1.52 | 0.95 | 12.81 | 18.8 | 10.37 | 1158 | <.001 | NA |
| effect change by layer 13 in BU | 14.99 | 1.52 | 0.95 | 11.99 | 17.98 | 9.83 | 1158 | <.001 | NA |
| effect change by layer 14 in BU | 13.49 | 1.52 | 0.95 | 10.5 | 16.48 | 8.85 | 1158 | <.001 | NA |
| effect change by layer 15 in BU | 11.41 | 1.52 | 0.95 | 8.42 | 14.4 | 7.48 | 1158 | <.001 | NA |
| effect change by layer 16 in BU | 8.9 | 1.52 | 0.95 | 5.9 | 11.89 | 5.83 | 1158 | <.001 | NA |
| effect change by layer 17 in BU | 6.22 | 1.52 | 0.95 | 3.23 | 9.21 | 4.08 | 1158 | <.001 | NA |
| effect change by layer 18 in BU | 3.67 | 1.52 | 0.95 | 0.67 | 6.66 | 2.4 | 1158 | 0.02 | NA |
| effect change by layer 19 in BU | 1.42 | 1.52 | 0.95 | -1.57 | 4.41 | 0.93 | 1158 | 0.35 | NA |
| effect change by layer 20 in BU | -0.4 | 1.52 | 0.95 | -3.4 | 2.59 | -0.27 | 1158 | 0.79 | NA |
| change of layer 2 effect by TD | -0.11 | 2.16 | 0.95 | -4.34 | 4.12 | -0.05 | 1158 | 0.96 | 0.98 |
| change of layer 3 effect by TD | -0.36 | 2.16 | 0.95 | -4.59 | 3.87 | -0.17 | 1158 | 0.87 | 0.98 |
| change of layer 4 effect by TD | -0.58 | 2.16 | 0.95 | -4.81 | 3.65 | -0.27 | 1158 | 0.79 | 0.98 |
| change of layer 5 effect by TD | -0.69 | 2.16 | 0.95 | -4.92 | 3.54 | -0.32 | 1158 | 0.75 | 0.98 |
| change of layer 6 effect by TD | -0.7 | 2.16 | 0.95 | -4.94 | 3.53 | -0.33 | 1158 | 0.74 | 0.98 |
| change of layer 7 effect by TD | -0.62 | 2.16 | 0.95 | -4.85 | 3.61 | -0.29 | 1158 | 0.77 | 0.98 |
| change of layer 8 effect by TD | -0.46 | 2.16 | 0.95 | -4.69 | 3.77 | -0.21 | 1158 | 0.83 | 0.98 |
| change of layer 9 effect by TD | -0.29 | 2.16 | 0.95 | -4.52 | 3.94 | -0.13 | 1158 | 0.89 | 0.98 |
| change of layer 10 effect by TD | -0.05 | 2.16 | 0.95 | -4.28 | 4.18 | -0.02 | 1158 | 0.98 | 0.98 |
| change of layer 11 effect by TD | 0.16 | 2.16 | 0.95 | -4.07 | 4.39 | 0.08 | 1158 | 0.94 | 0.98 |
| change of layer 12 effect by TD | 0.3 | 2.16 | 0.95 | -3.93 | 4.54 | 0.14 | 1158 | 0.89 | 0.98 |
| change of layer 13 effect by TD | 0.31 | 2.16 | 0.95 | -3.92 | 4.54 | 0.14 | 1158 | 0.89 | 0.98 |
| change of layer 14 effect by TD | 0.32 | 2.16 | 0.95 | -3.91 | 4.55 | 0.15 | 1158 | 0.88 | 0.98 |
| change of layer 15 effect by TD | 0.45 | 2.16 | 0.95 | -3.78 | 4.68 | 0.21 | 1158 | 0.83 | 0.98 |
| change of layer 16 effect by TD | 0.79 | 2.16 | 0.95 | -3.44 | 5.02 | 0.37 | 1158 | 0.71 | 0.98 |
| change of layer 17 effect by TD | 1.26 | 2.16 | 0.95 | -2.97 | 5.49 | 0.59 | 1158 | 0.56 | 0.98 |
| change of layer 18 effect by TD | 1.75 | 2.16 | 0.95 | -2.48 | 5.98 | 0.81 | 1158 | 0.42 | 0.98 |
| change of layer 19 effect by TD | 2.22 | 2.16 | 0.95 | -2.01 | 6.45 | 1.03 | 1158 | 0.3 | 0.98 |
| change of layer 20 effect by TD | 2.58 | 2.16 | 0.95 | -1.65 | 6.81 | 1.2 | 1158 | 0.23 | 0.98 |

*Supplementary table 13 Robust linear mixed-effects model parameter estimates for right posterior insula.*
